# Discovery of metabolites produced by reactions between central carbon metabolites and cysteine that mark inflammatory macrophages

**DOI:** 10.64898/2026.03.18.712640

**Authors:** Nicholas L. Arp, Felicia Deng, Jorgo Lika, Gretchen L. Seim, Paulo Falco Cobra, Carlos Mellado Fritz, Steven V. John, Swetha Rathinaraj, Bridget E. Shields, Daniel Amador-Noguez, Katherine Henzler-Wildman, Jing Fan

## Abstract

Identifying metabolites and metabolic reactions specific to a cellular state, such as inflammatory state in immune cells, is of great interest, as it can provide important biomarkers and point to compounds and reactions of specific biological functions. However, many cell state-specific metabolites remain in the unannotated part of metabolome. Here we identified a series of sulfur-containing metabolites that are actively produced in macrophages upon classical activation, but not in resting state or alternative activation state. Isotopic tracing, in vitro assays and genetic perturbations further revealed that they are formed from reactions between free cysteine and several important intermediates in glycolysis and TCA cycle. Upon classical activation, macrophages specifically upregulate the import of cystine via *Slc7a11*, supporting the production of these adducts. Their production dynamically responds to changes in central metabolism, environmental nutrient levels, and is regulated by nitric oxide. Finally, we confirmed these newly identified compounds also present in human samples, and most of them are significantly elevated in inflammatory granuloma annulare lesions. This work elucidated a previously uncharted part of metabolic network that is associated with inflammation and metabolic stress condition, which has important implications and set foundation for many future discoveries.

## INTRODUCTION

Uncovering new metabolites and biochemical reactions has always been a foundational question in metabolism research. However, much of the mammalian metabolome remains unknown to this day. Especially, as decades of efforts have been largely concentrated on mapping out the central “ubiquitous” pathways in the mammalian metabolic network, the metabolites and reaction that are uniquely present in, and important for, certain biological states is a great knowledge gap.

Identifying metabolites associated with specific cellular states is of great significance, as it not only provides useful markers, but also opens the door to a series of discoveries elucidating the dynamic regulation and the biological functions. Particularly, there has been great interest in elucidating the metabolites and metabolic reactions characteristic to different immune polarization states in macrophages. In response to different signals associated with infection, inflammation, or injury, macrophages can take on a diverse spectrum of functional polarization states, with the inflammatory classical activation state (“M1”,commonly induced by lipopolysaccharide and interferon-γ) and the anti-inflammatory alternative activation state (“M2’,commonly induced by IL-4 and IL-13) on two sides of the spectrum. The best-known example of a metabolite that is specifically associated with macrophage polarization is itaconate, “the poster child of immunometabolism” ^1,2^. Itaconate were found to be a mammalian metabolite 15 years ago (prior to that it was known as a microbial metabolite), whose production by macrophages is specifically upregulated in classical activation state^3^. Since then, a great wave of research has identified the reactions responsible for the production and catabolism of itaconate in mammalian cells^4,5^, elucidated the mechanisms regulating the rise-and-fall of itaconate level during immune response^6,7^, and revealed the many functional impacts of itaconate and the mechanisms mediating these impacts^2^. Not only can itaconate act as an anti-microbial compound released by classical activated macrophages^8,9^, intracellular itaconate was found to be crucial for orchestrating the dynamic immune response by modifying key signaling proteins including KEAP1, ATF3, NRLP3^10–12^, and regulating other metabolic enzymes such as GAPDH and SDH^13–15^.

It is reasonable to expect there are more metabolites like itaconate that are specific to an immune activation state. Activated macrophages perform many important functions that are not active in the resting state, such as the production of reactive oxygen and nitrogen species for pathogen killing, phagocytosis, and production of cytokines. The activation of these functions poses associated metabolic demands and metabolic stress. Metabolites and metabolic reactions are important for supporting immune functions, adapting to metabolic stress, or regulating immune response are expected to be upregulated. Indeed, growing research shows that macrophages undergo substantial and dynamic reprogramming throughout the broad metabolic network upon activation. However, studies thus far have been limited to the metabolic remodeling in the known annotated part of metabolic network. There are likely other interesting “polarization state-specific” metabolites hidden in the uncharted part of metabolic network, just like how itaconate was to us 15 years ago. Identifying these metabolites is not trivial. Most widely applied metabolomic workflows depend on database matching, for instance, searching the exact *m/z* and MS/MS pattern in an experimental mass spectrometry dataset against a comprehensive metabolite database, such as KEGG and the human metabolome database (HMDB)^16,17^. However, metabolites that are truly uniquely produced in specific conditions are likely never described before. Moreover, unlike proteomics, where the similar structure of peptides makes it highly reliable to generate expected MS/MS spectra based on the amino acid sequence, small metabolites can be very diverse in chemical structure, thus there is no consistent template ^18^. These challenges can make interesting discoveries hidden in the “dark” parts of metabolic network.

Here we searched for previously unknown metabolites that are uniquely produced in large quantities only in classically activated macrophages, using an integrated workflow. We started with LCMS-based discovery metabolomics coupled with statistical analyses to extract interesting features and identify formulas, followed by symmetric isotopic tracing and *in vitro* experiments to elucidate the reaction producing these metabolites. NMR analysis was applied in addition to LCMS to elucidate the structures. Genetic and pharmacological perturbations of upstream enzymes and nutritional perturbations further confirmed their metabolic sources. Together, we identified a group of previously unknown metabolites that are formed from the reaction between cysteine and several key metabolites from glycolysis and TCA cycle, including most notably, glyceraldehyde-3-phosphate (GAP), dihydroxyacetone phosphate (DHAP), and α-ketoglutarate (αKG). Upon classical activation, macrophages greatly upregulate glycolysis and dynamically remodel TCA cycle, causing the fluctuation or accumulation of these intermediates^6,7,19–21^. The upregulation in these cysteine adducts on one hand responds to the dynamic changes in central metabolism, while on the other hand, is enabled by the active upregulation of cystine transporter SLC7A11 and unique increase of cysteine level among all amino acids upon classical activation. Nitric oxide (NO), a bioactive molecule produced by activated macrophages, is also a regulator for the production of these newly identified compounds. These metabolites specifically mark classically activated macrophages and are significantly increased in patient samples of a macrophage-driven inflammatory disease, granuloma annulare.

## RESULTS

### Discovery metabolomics identified a group of sulfur-containing compounds specifically accumulate in classically activated macrophages

To systematically identify metabolites that are uniquely associated with classical activation state, we performed LCMS-based metabolomics analysis of bone marrow derived macrophages (BMDM) that are unstimulated or stimulated with lipopolysaccharide (LPS) and interferon-γ (IFNγ) and extracted all features (LCMS peaks). Statistical analysis was performed to identify meaningful metabolomic features specific to the activated state, using the criteria that the feature is present at significant amount in stimulated macrophages, but are not detected or substantially less abundant (> 2.5-fold change, p <0.05) in unstimulated macrophages (Fig 1A). A set of LCMS features corresponding to the best-known metabolic markers of classical activation, citrulline (including its isotopic peaks and MS adducts) were captured among the top hits identified by this analysis, providing a positive control. This analysis also identified many classical activation-specific features whose exact *m/z* do not match any known metabolite HMDB database, suggesting possible new mammalian metabolites to be discovered.

**Figure 1.**
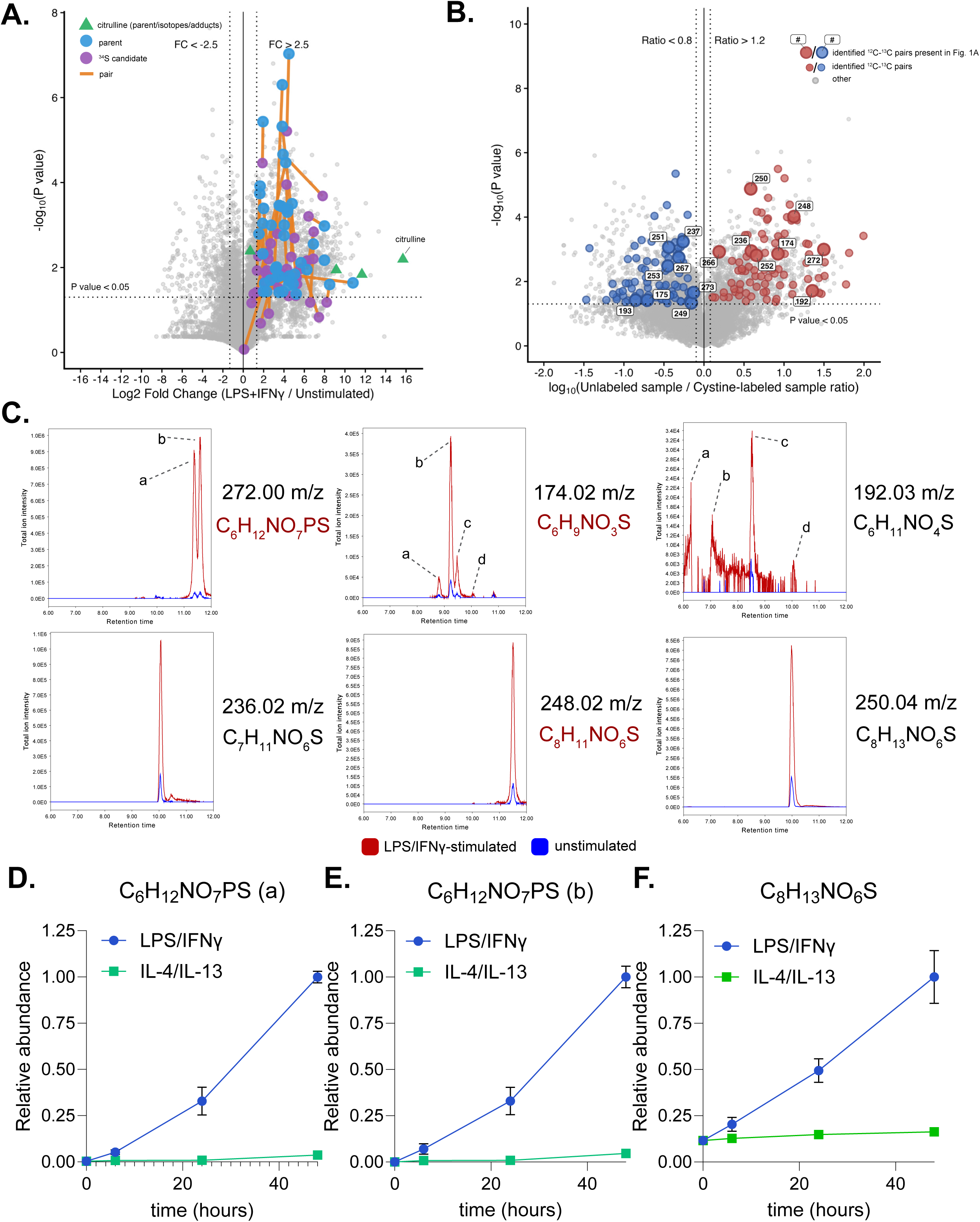
Discovery of cysteine adducts in classically activated macrophages. **A.** volcano plot of discovery metabolomics comparing BMDMs that stimulated with LPS/IFNγ (at 48h post stimulation) versus unstimulated. Colored points indicate parent feature (blue circle) and ^34^S isotope peak candidates (purple circle) with orange line indicated parent-isotope pair. Only parent feature that meet > 2.5-fold change and significance < 0.05 are labeled. Citrulline (green triangle) (and its isotope peaks and MS adducts) are labeled as a positive control for classical activation induced metabolites. Each point represents the mean of n = 3 biological replicates. **B.** volcano plot of metabolomic features in parallel cells cultured in media containing unlabeled cystine versus 3,3-^13^C_2_-L-cystine. Pairs with significant isotopic shift are colored blue and red. Pairs that match parent features in (A) are labeled with m/z values. Each point represents the mean, n = 3 biological replicates. **C.** Extracted ion chromatograms (EICs) for major cysteine adducts in LPS/IFNγ-stimulated BMDM (red) compared to in unstimulated BMDMs (blue). Identified molecular formulas based on isotopic distribution and exact mass are labeled. Formulas shown in red do not correspond to known mammalian metabolites. **D-F.** The changes in major cysteine-adducts in BMDMs over a time course of LPS/IFNγ stimulation (blue) versus IL-4/IL-13 stimulation (green). Mean +/- standard deviation, n= 3 biological replicates.

To identify these metabolites, we first grouped the features to those that originate from the same compound (as isotopic peaks and MS adducts), based on their retention time, covariance, and mass intervals. This revealed the natural isotopic distribution of these compounds. Interesting, many of the high abundance classical activation-specific metabolites contain a S^34^ isotopic peak (+1.996), at approximately 4% of the signal of the base peak (Fig 1A). This is highly indicative that these compounds contain one sulfur atom. Similarly, we analyzed the relative abundance of other isotopic peaks (primarily ^13^C at +1.0034 from the base peak). The isotopic distribution together with exact mass and chemical feasibility allowed us to hypothesize the chemical formulas of many metabolites.

As cysteine is a main sulfur source in these cells, we hypothesized that the sulfur-containing compounds may be derived from cysteine. To test this, we performed isotopic tracing with 3,3-^13^C_2_-L-cystine, because cystine in the media is the major source of cellular cysteine. Macrophages cultured in media containing either unlabeled or isotope-labeled L-cystine for 24 hours were analyzed by LC-MS. Cysteine-derived metabolites are expected to have a +1.0034 Da mass shift in labeled samples compared to unlabeled. Analysis of the characteristic ^12^C-^13^C isotopologue pairs in this dataset broadly capture cysteine-derived metabolome (Fig. 1B). Indeed, many of the sulfur-containing metabolites that significantly accumulate upon classical activation are derived from cystine. The formulas of these metabolites were identified as: C_6_H_12_NO_7_PS (272 m/z), C_8_H_13_NO_6_S (250 m/z), C_8_H_11_NO_6_S (248 m/z), C_6_H_9_NO_3_S (174 m/z), C_7_H_11_NO_6_S (236 m/z), and C_6_H_11_NO_4_S (192 m/z). Some of these formulas correspond to multiple peaks of the same m/z but retain differently on liquid chromatography, suggesting there are chemically distinct isomers. All these isomers similarly show substantial accumulation upon macrophage classical activation (Fig. 1C).

To understand whether the accumulation of these metabolites in macrophages is specific to classical activation, we then compared the dynamic changes of these newly identified compounds over the time courses of classical activation induced by LPS+ IFNγ versus alternative activation induced by IL-4 and IL-13, focusing on the three most abundant species: two isomers of C_6_H_12_NO_7_PS and C_8_H_13_NO_6_S. Compared to their strong and continuous accumulation over 48 hours of classical activation time course, their change over alternative activation are minimal (Fig. 1D-F). Thus, these newly identified compounds are specific metabolic features that distinguish classically activated macrophages from resting macrophages or alternatively activated macrophages.

### Isotopic labeling revealed these compounds are derived from central carbon metabolites and cysteine

To understand the additional metabolic sources of these compounds, we applied isotopic tracing approach using the tracers including D-glucose and L-glutamine, which are the two major carbon sources, alongside the previously described, L-cystine, which is a major source of sulfur. Cells were cultured in a group of parallel conditions using culture media that are identical in chemical composition but contain different isotopes of designated nutrients for 24 hours (Fig. 2A and B). As expected, within 24 hours, labeling with 3,3’-^13^C_2_-L-cystine resulted in ∼80% of intracellular cysteine being labeled (Fig. 2C), and labeling with U-^13^C-L-glutamine or U-^13^C-D-glucose resulted in most intracellular glutamine or glycolytic intermediates being labeled, respectively (Fig. 2D and E). As demonstrated in Figure 1B, all the classical activation-specific sulfur-containing compounds we identified above get 1-labeled from 3,3’-^13^C_2_-L-cystine (Fig. 2F-I), validating that cysteine contributes to parts of these metabolites.

**Figure 2.**
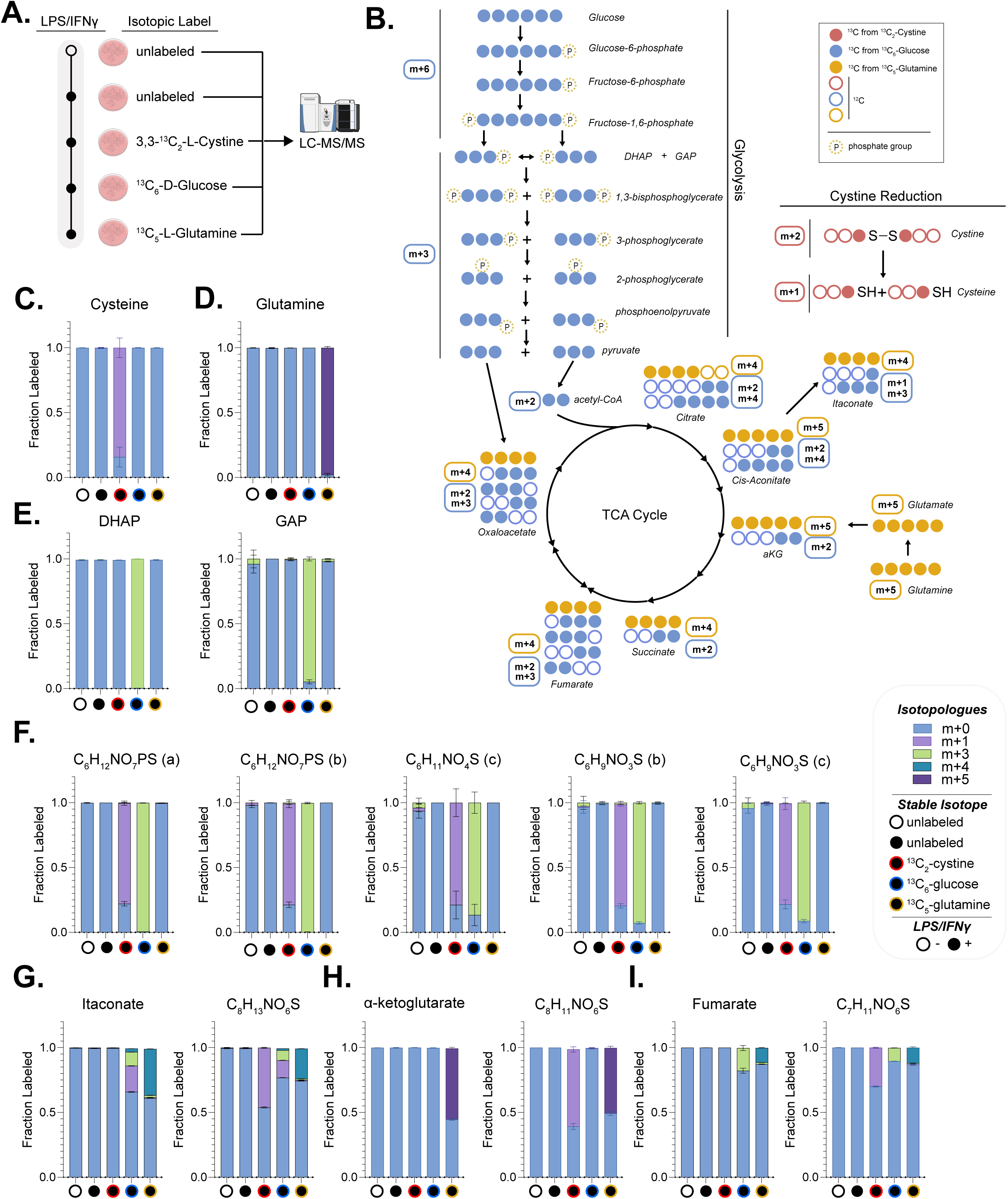
Isotope tracing reveals the metabolic sources of cysteine adducts. **A.** Experimental design for symmetric 24-hour stable isotope tracing using U-^13^C-D-glucose, 3,3-^13^C_2_-L-cystine, U-^13^C_5_-L-glutamine, or unlabeled media control in LPS/IFNγ-stimulated BMDMs (48h post stimulation). Unstimulated BMDMs cultured in unlabeled media were included as an additional control condition. **B.** Metabolic pathway schematic showing major expected isotopologue patterns (m+0 through m+6) for glycolysis and TCA cycle intermediates with each tracer. Legend indicates carbon source by color. **C-E.** Fractional labeling of (**C**) cysteine, (**D**) glutamine, and (**E**) glycolytic intermediates DHAP and GAP showing labeling incorporation from each tracer. **F.** Fractional labeling of C_6_H_12_NO_7_PS, C_6_H_11_NO_3_S, and C_6_H_9_NO_3_S (the high abundance isomers shown for each formula) show predominant 3 carbon labeling from glucose, matching lower glycolytic intermediates. **G-I.** Similar labeling pattern of (**G**) itaconate and C_8_H_13_NO_6_S; (**H**) C_8_H_11_NO_6_S and α-ketoglutarate; (**I**) fumarate and C_7_H_11_NO_6_S, from glucose and glutamine.

Among these metabolites, all of those containing six carbon, including C_6_H_12_NO_7_PS (two isomers), C_6_H_9_NO_3_S (two isomers), and C_6_H_11_NO_4_S (at least one major isomer) become nearly 100% 3-labeled from U-^13^C-glucose, but do not label from U-^13^C-glutamine (Fig. 2F), following the labeling pattern of 3-carbon lower glycolytic intermediates like glyceraldehyde-3-phosphate (GAP) and dihydroxyacetone phosphate (DHAP) (Fig. 2E). These results lead to the hypothesis that these 6-carbon compounds resulted from the reaction between cysteine (providing 3 carbon and one sulfur and one nitrogen), and a 3-carbon glycolytic intermediate.

The other subgroup of these newly identified metabolites, those containing 7 to 8 carbons (C_8_H_13_NO_6_S, C_8_H_11_NO_6_S, and C_7_H_11_NO_6_S), all get significantly labeled from U-^13^C-L-glutamine, a major carbon source of the TCA cycle, leading to the hypothesis that these compounds resulted from the reaction between cysteine and certain compounds derived from the TCA cycle. Indeed, we found the labeling patten of C_8_H_13_NO_6_S, C_8_H_11_NO_6_S, and C_7_H_11_NO_6_S, from both U-^13^C-glucose and U-^13^C-glutamine, follow the labeling pattern of itaconate, α-ketoglutarate (αKG), and fumarate, respectively (Fig. 2G-I).

Next, we examined the turnover of these compounds in classically activated macrophages by kinetic glucose tracing (Fig. 3A). U-^13^C-glucose fully labeled glycolytic intermediates, such as GAP and DHAP, within 30 minutes. The 6-carbon sulfur-containing compounds get 3-labeled gradually. Both isoforms of C_6_H_12_NO_7_PS become ∼55% labeled by 30 minutes and nearly completely labeled by 4 hours (Fig. 3B). Labeling rate of C_6_H_11_NO_4_S differs by isomers, with the labeling half time ranging from ∼30 minutes to 1 hour (Fig. 3B). Overall, the delay of their labeling rate compared to the glycolytic intermediates suggests they are downstream, and the relatively short labeling half-life (< 1 hour) indicates they are actively turned over in cells.

**Figure 3.**
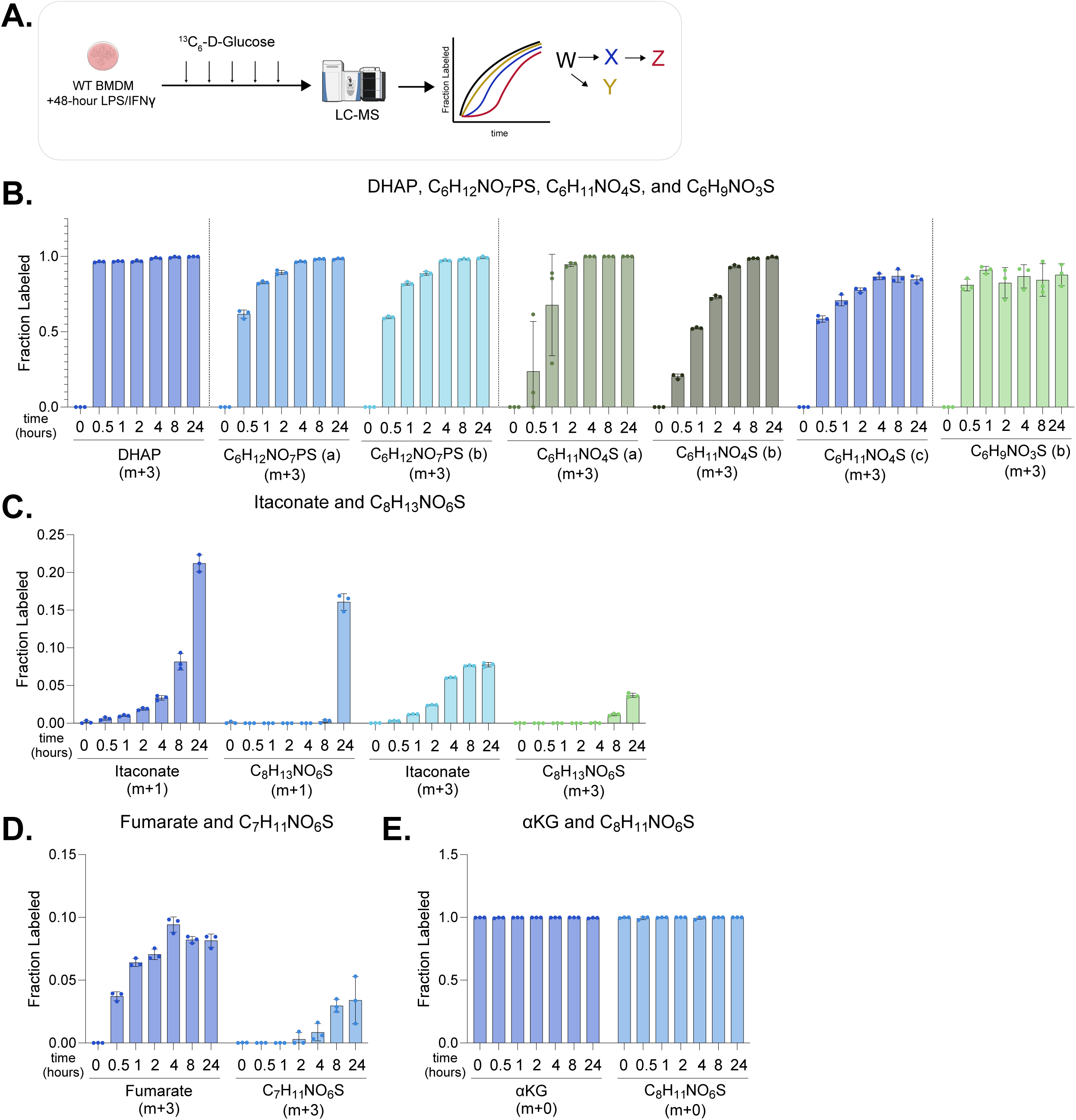
Kinetic labeling of cysteine adducts. **A.** Experimental design for time-resolved U-^13^C-D-glucose tracing in LPS/IFNγ-stimulated BMDMs. **B.** Time course showing m+3 labeled fraction in DHAP and glycolysis-derived cysteine adducts C_6_H_12_NO_7_PS (a/b), C_6_H_11_NO_4_S (a/b/c), and C_6_H_9_SO_3_S (b). Mean +/- standard deviation (SD), n = 3 biological replicates. **C.** Fractional labeling (showing the major isotopologues, m+1 and m+3) of itaconate and C_8_H_13_NO_6_S showing accumulation of labeled species over time. Mean +/- SD, n = 3 biological replicates. **D.** Fractional labeling (showing the major isotopologue, m+3) of fumarate and C_7_H_11_NO_6_S over time. **E.** Fractional labeling of α-ketoglutarate (m+0) and C_8_H_11_NO_6_S (m+0) showing these compounds remain largely unlabeled from glucose in stimulated BMDMs. Mean +/- SD, n = 3 biological replicates.

Similarly, the labeling of C_8_H_13_NO_6_S and C_7_H_11_NO_6_S shows similar patterns that track with itaconate and fumarate, respectively, but slower in rates (Fig. 3C-D), placing them downstream of these TCA cycle metabolites. C_8_H_11_NO_6_S does not get significantly labeled from glucose, mirroring that αKG does not get significantly labeled from glucose in classically activated macrophages (Fig. 3E). The delay of C_7_H_11_NO_6_S labeling compared to fumarate labeling is the most notable (Fig. 3D). This is consistent with the observation that C_7_H_11_NO_6_S only become ∼30% labeled from labeled cystine by 24 hours (Fig. 2H), suggesting it is the compound of the slowest turnover rate within this group.

### In vitro confirmation of biochemical reactions producing these metabolites

Based on the tracing experiments above, we hypothesized these classical activation-specific sulfur-containing metabolites are produced from reactions between free cysteine and certain metabolites derived from lower glycolysis and TCA cycle. Specifically, the formula C_6_H_12_NO_7_PS fits the dehydration condensation between cysteine and GAP or DHAP. C_6_H_11_NO_4_S and C_6_H_9_NO_3_S can be also produced in related reactions, as they only differ from C_6_H_12_NO_7_PS by a HPO3 (dephosphorylation) or H3PO4 group, respectively. Methylglyoxal (MGO), a by-product of glycolysis downstream of DHAP, may also be responsible for generating some of these compounds in cells, especially given a recent report nicely demonstrated it can react with cysteine non-enzymatically and produce lactoyl-cysteine (some isomers of C_6_H_11_NO_4_S)^22^. Regarding the subgroup appearing to be downstream of TCA cycle: C_8_H_13_NO_6_S and C_7_H_11_NO_6_S show labeling patterns following that of itaconate and fumarate, respectively, and the formulas fit the addition products of itaconate or fumarate with cysteine, and C_8_H_11_NO_6_S showed labeling patterns following that of αKG and fits the formula of dehydration condensation of cysteine and αKG. We therefore tested the hypothesis the reaction between cysteine and itaconate, fumarate, and αKG generate these observed metabolites respectively.

Each of the hypothesized precursors in glycolysis or TCA cycle was co-incubated with free cysteine at concentrations selected to approximate reported intracellular levels^5,23–25^ (specified in Fig. 4A), in neutral buffer environment at 37°C for 1 or 24 hours. We found all these hypothesized metabolites can react with cysteine and generated a series of products that collectively covered all the classical activation-induced cysteine adducts observed in cells (Fig. 4B-G), with matching retention times and MS/MS spectra (Supp. Fig. 1-6). The reactions include two general types. First, itaconate and fumarate react with cysteine through addition reaction, generating C_8_H_13_NO_6_S and C_7_H_11_NO_6_S matching the corresponding LCMS features observed in stimulated cells (Fig. 4B-C) with MS/MS fragments correspond to cysteine and methylsuccinyl-group or succinyl-group respectively (Supp. Fig. 1A-C and 2A-C). Furthermore, these fragments carry expected label from 3,3’-^13^C_2_-L-cystine, U-^13^C-D-glucose and U-^13^C-L-glutamine, confirming their assignments (Supp. Fig. 1C-F and 2C-F). These results are consistent with previous literature showing S-itaconylation and S-succinylation can non-enzymatically occur on cysteine residual of proteins (or free cysteine)^10,12,26,27^, provide strong confidence in identifying them as the S-itaconyl-cysteine (Ita-Cys) and S-succinyl-cysteine (Suc-Cys) (Supp. Fig. 1G and 2G). The second group of reactions appear to occur between cysteine and carbonyl group. The reaction of GAP and DHAP both produce various products of the formulas C_6_H_12_NO_7_PS, C_6_H_11_NO_4_S and C_6_H_9_NO_3_S, with different isomer distributions. Most notably, GAP only produces isomer (a) of C_6_H_12_NO_7_PS observed in cells, while DHAP produces both (Fig. 4D-E, Supp. Fig. 3A-G, Supp. Fig. 4A-G, and Supp. Fig. 5A-I). C_6_H_11_NO_4_S and C_6_H_9_NO_3_S can also be produced from MGO reacting with cysteine. Based on a recent study ^22^, some of the three detected isomers of C_6_H_11_NO_4_S resulted from this reaction could be lactoyl-cysteine (Fig. 4F and Supp. Fig. 4A-F). The reaction of cysteine with αKG produced dehydration condensation products. The product of αKG shows LC retention and MS/MS matches C_8_H_11_NO_6_S (indicated as αKG-Cys thereafter) observed in cells (Fig. 4G and Supp Fig 6A-C).

**Figure 4.**
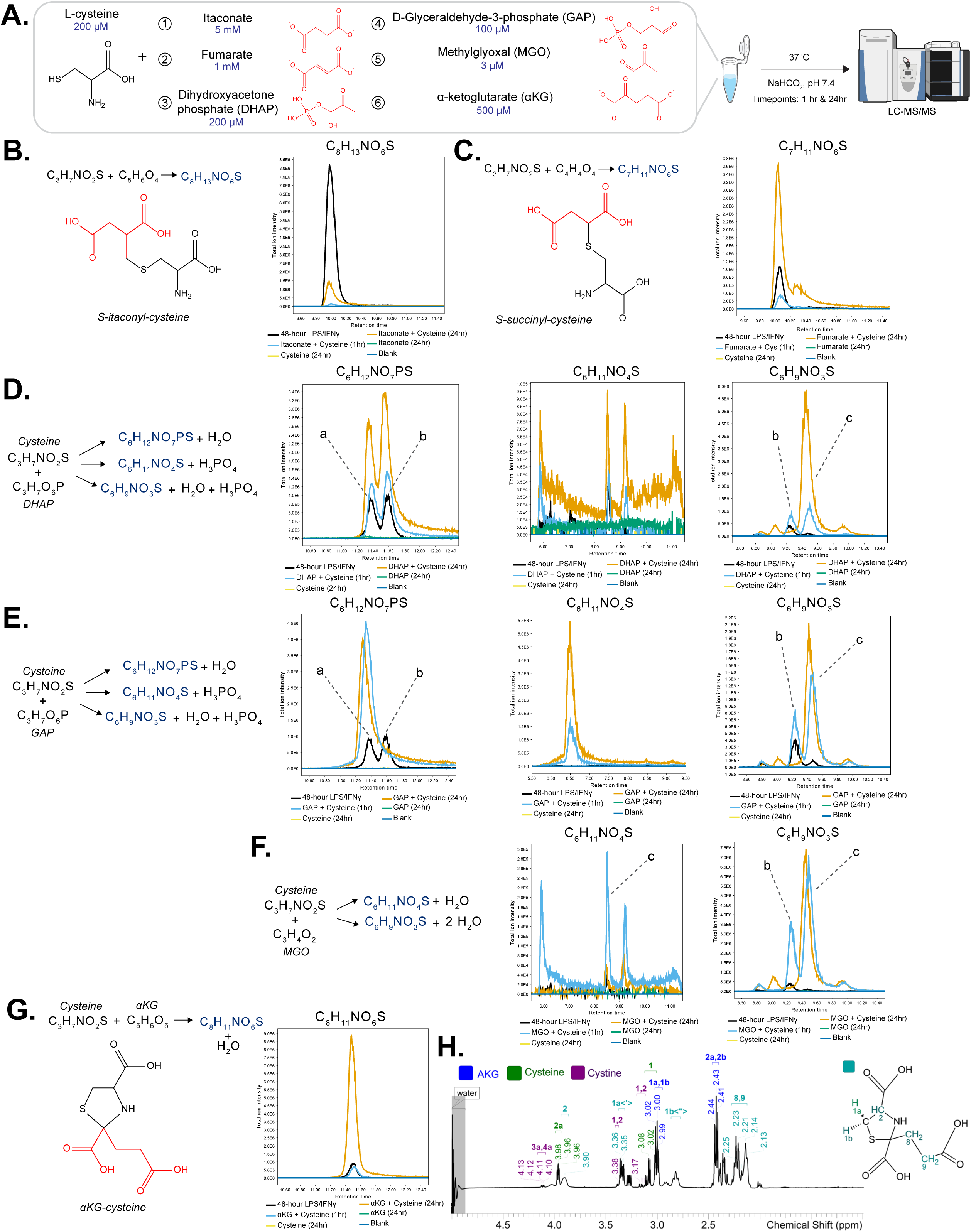
Reactions produce central carbon metabolite-cysteine adducts. **A.** Experimental design of the in vitro assays. Cysteine (200 μM) was incubated with indicated metabolites at indicated concentrations (chosen to match physiological range) in NaHCO_3_ buffer (pH 7.4, 37 °C) for 1 or 24 hours. Products were analyzed by LC-MS/MS. **B-G.** Left: the reactions of cysteine with (**B**) itaconate, (**C**) fumarate, (**D**) DHAP, (**E**) GAP, (**F**) methylglyoxal, and (**G**) α-ketoglutarate, with the formula of various detected products indicated. Right: EICs showing the formation of each of the products at 1h and 24h co-incubation, compared to blank or signal substrate control. For comparison, the EIC of these compounds detected in stimulated macrophages (shown in black) were overlaid onto the EIC of these products formed in vitro. **H.** ^1^H NMR spectrum of the reaction mix of α-ketoglutarate co-incubating with cysteine (room temperature, 24 hours; NaHCO_3_, pH 7.4), confirming the thiazolidine containing structure of the product (shown in right). Peak annotations are shown on the spectrum: resonances corresponding to α-ketoglutarate (blue), cysteine (green), cystine (purple), and reaction product (cyan).

**Figure 5.**
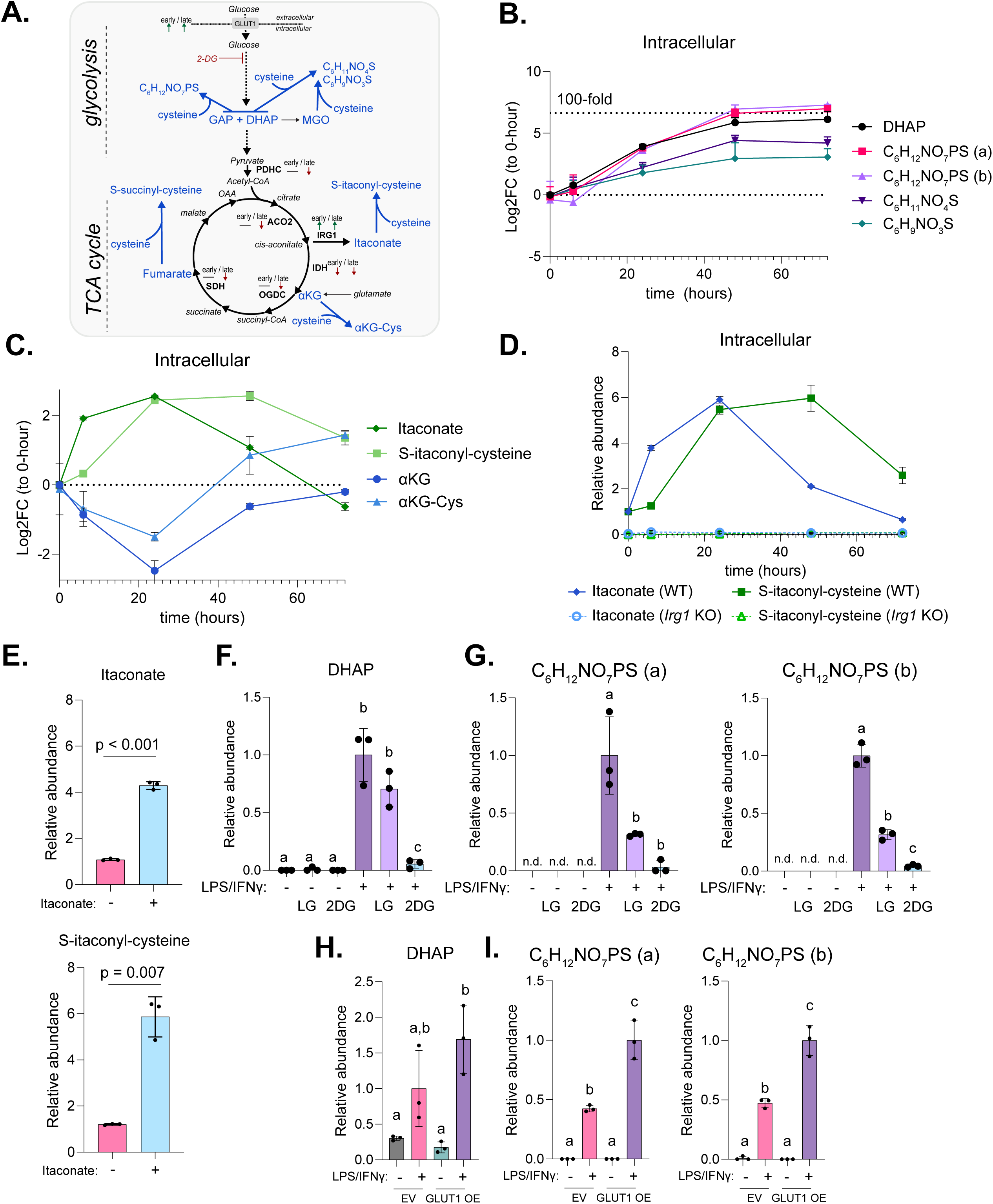
Cysteine adducts respond to metabolic perturbations in central carbon metabolism. **A** Schematics showing the dynamic remodeling of central carbon metabolism upon LPS/IFNγ-stimulation, and by the experimental perturbations of key steps **B-C.** Dynamic changes in glycolysis-derived and TCA cycle-derived cysteine adducts in macrophages, over a time course upon LPS/IFNγ-stimulation. Relative abundance at each time point shown as Log_2_ fold change relative to in unstimulated macrophages (0h). The relative level of their corresponding precursors, DHAP, α-ketoglutarate, and itaconate were also shown for comparison.. **D.** Relative abundance of itaconate and Ita-Cys (C_8_H_13_NO_6_S) in wildtype (WT) versus *Irg1* KO BMDMs over the time course of LPS/IFNγ-stimulation. **E.** Itaconate supplementation (1 mM) increases Ita-Cys level in unstimulated WT BMDMs. **F-G.** Glycolysis inhibition by 2-deoxy-glucose (2DG, 1 mM) or culture cells in low glucose (LG, 1 mM) reduces DHAP and C_6_H_12_NO_7_PS accumulation in LPS/IFNγ-stimulated BMDMs. **H-I.** GLUT1 overexpression in RAW264.7 cells increases glycolysis-derived cysteine adducts accumulation upon LPS/IFNγ-stimulation (48h time point). **(B-I)** Mean +/-standard deviation (SD), n = 3 biological replicates. Statistical analysis: (**E**) unpaired two-tailed t-test, or (**F-I**) one-way ANOVA with Tukey’s post-hoc test (bar with different letters indicate significant differences, p< 0.05).

**Figure 6.**
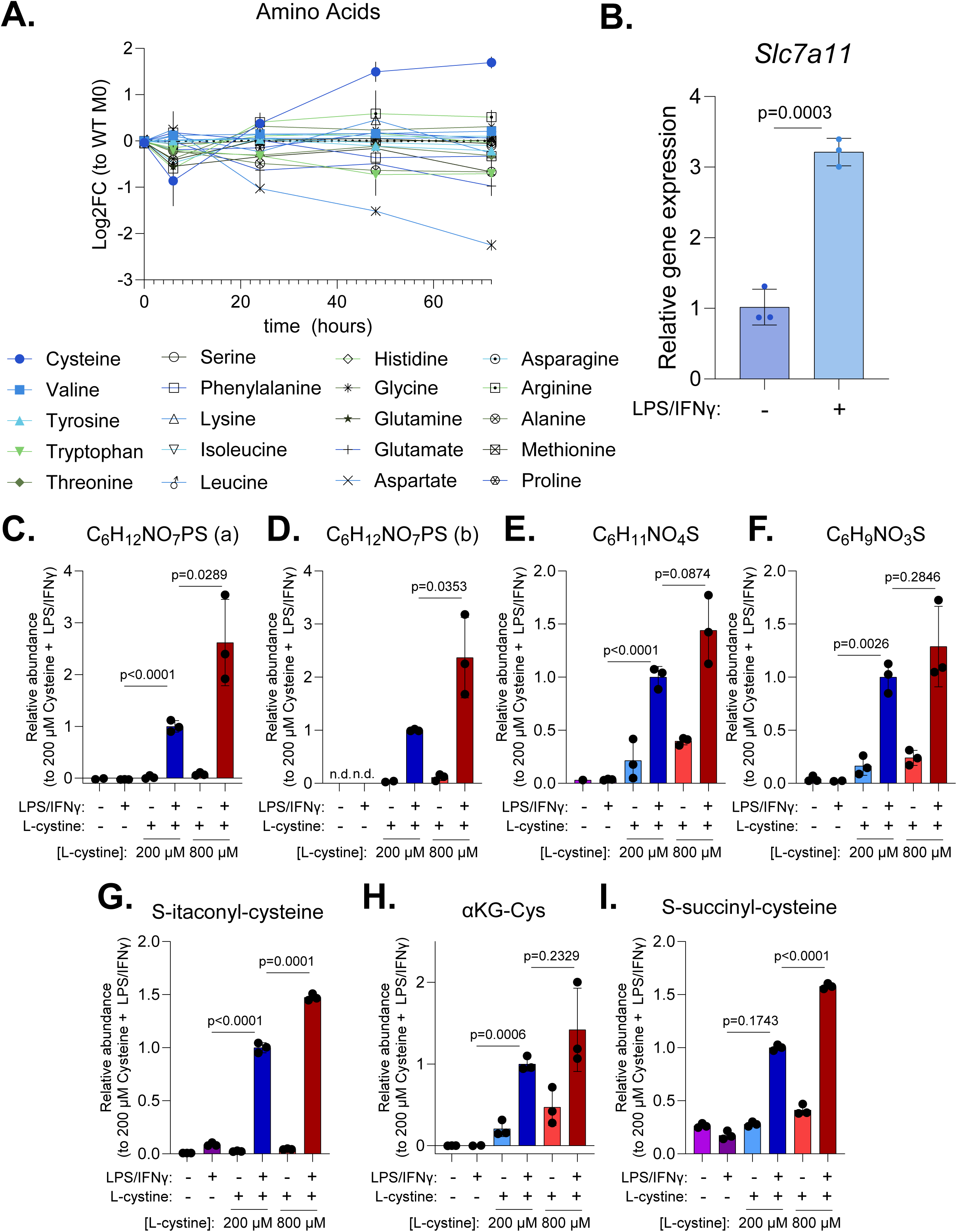
Cystine uptake and availability influence cysteine adducts formation. **A.** Relative intracellular abundance of all 20 amino acids in BMDMs over a 72-hour time course upon LPS/IFNγ-stimulation. Relative abundances are shown as Log2 fold change relative to 0h time point (unstimulated BMDM). Mean +/- standard deviation (SD), n = 3 biological replicates. **B.** *Slc7a11* transcript increases with LPS/IFNγ-stimulation in BMDMs. Mean +/- SD, n = 3 biological replicates. Statistical comparison by unpaired two-tailed t-test with exact p-value reported. **C-I.** Relative abundances of cysteine adducts in BMDMs cultured in media with standard cystine concentration (200 μM), high cystine (800 μM), or no cystine (0 μM) with or without LPS/IFNγ-stimulation for 48 hours. n.d. = not detected. Mean +/- SD, n = 3 biological replicates. Statistical comparisons were performed between stimulated high cystine or no cysteine conditions compared to standard cystine concentrations using unpaired two-tailed t-test with exact p-value reported.

Examining the rates of these reactions, several interesting observations emerged: (1) The reactions are generally faster for GAP, DHAP, MGO and αKG (the carbonyl containing substates) than fumarate and itaconate (the addition reactions), consistent with the observations in cell in the kinetic tracing experiments; (2) Some of the products continue to accumulate over the 24-hour time course, while others first increase then decrease, suggesting that they are converted or re-arranged to other more stable products. A notable example is C_6_H_9_NO_3_S produced from the reaction between cysteine and GAP, DHAP or MGO, isomer (b) first increase then decrease, replaced by isomer (c). Interestingly, isomer (b) is the major isomer observed in activated macrophages, suggesting the possibility that enzymatic activity producing or turnover some of the isomers in cells causing this distribution.

Overall, these results demonstrate that cysteine adducts that mark classically activated macrophages can be produced by non-enzymatic reactions with key metabolites in central metabolism (though it does not exclude enzymatic production). Particularly of interest, the significant reaction of GAP, DHAP, and αKG with cysteine, and their corresponding products, have never been reported before to the best of our knowledge. To better characterize the structure of the products, we next performed NMR analysis. NMR results demonstrate the reaction between αKG and cysteine primarily produce one cyclic adduct as shown in Figure 4H. The experimentally acquired ^1^H NMR data of the reaction mixture (Fig. 4H, in which resonances attributed to the reaction product are annotated in cyan, and resonances corresponding to αKG in blue, cysteine in green) match well with the theoretical chemical shifts of this product. This assignment is further corroborated by complementary 2D and heteronuclear NMR analyses, including ^13^C, ^13^C DEPT-135, ^1^H-^1^H COSY, and ^1^H-^13^C HSQC spectra (Supp. Fig. 7A-B). We were unable to confidently solve the structures of the products from reactions between GAP or DHAP with cysteine by NMR, due to the great complexity of the reaction (an evolving mixture including multiple isomers of C_6_H_12_NO_7_PS, C_6_H_11_NO_4_S, and C_6_H_9_NO_3_S are present, crowding the NMR spectra). Some possible products based on the chemical reactivity between identified substrates and LCMS information are provided in Supplementary Figure 8.

**Figure 7.**
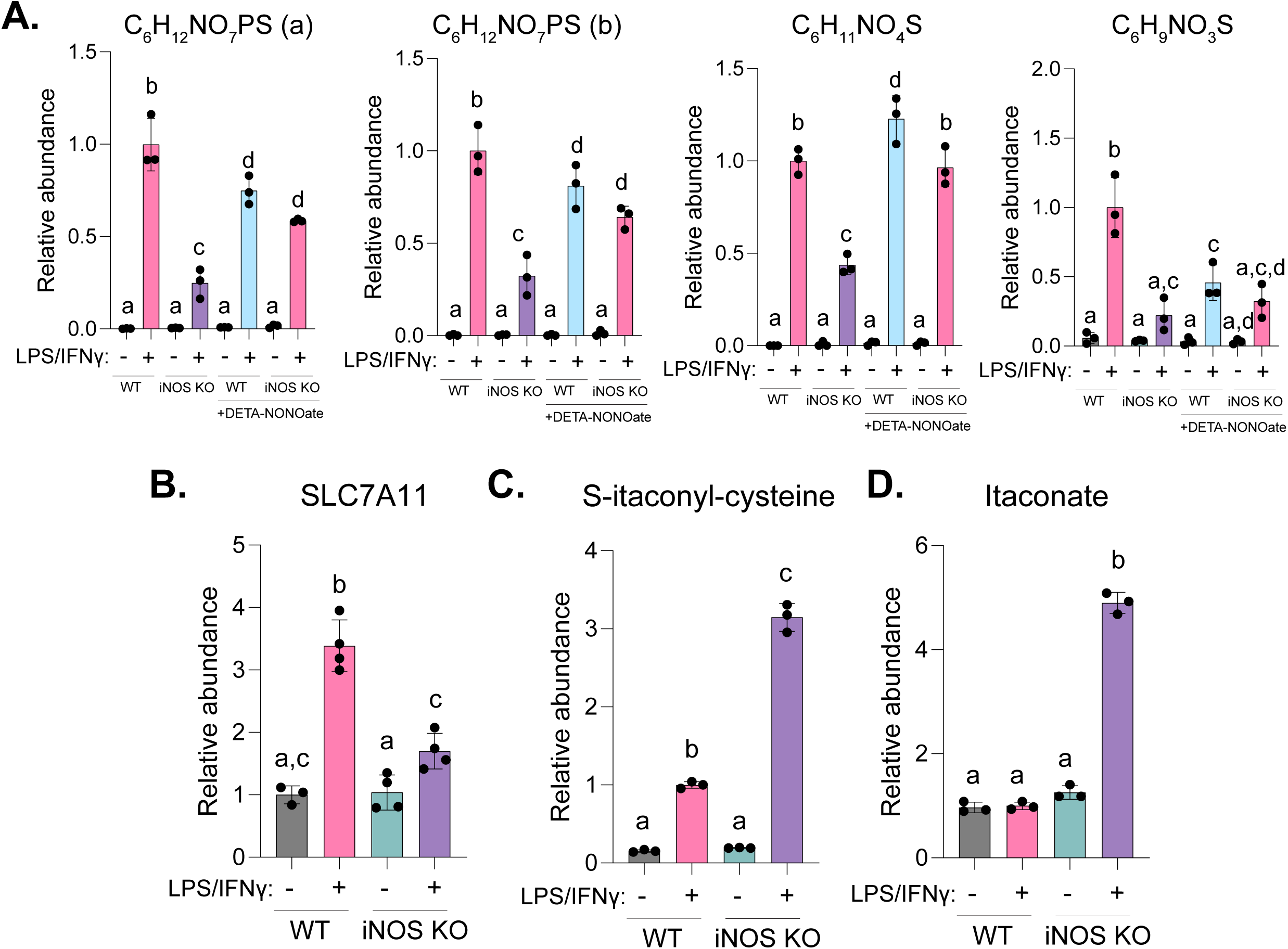
Nitric oxide regulates cysteine adducts. **A.** Relative abundance of glycolysis-derived cysteine adducts in wild-type (WT) versus iNOS KO BMDMs with or without LPS/IFNγ stimulation, with or without DETA-NONOate treatment (200 μM) for 48h. Mean +/- standard deviation (SD), n = 3 biological replicates. **B.** Relative SLC7A11 protein abundance from our recently deposited DIA proteomics dataset ^33^ (MassIVE accession MSV000100331) in WT and iNOS KO BMDMs with or without 48-hour LPS/IFNγ stimulation. Mean ± SD, n = 4 biological replicates (WT unstimulated: n = 3 after exclusion of one outlier by Grubbs’ test). **C-D.** Relative (C) Ita-Cys and (D) itaconate levels in WT and iNOS KO BMDMs with or without 48-hour LPS/IFNγ stimulation. Mean +/- SD, n = 3 biological replicates. (**A-D**) Bar with different letters indicate significant differences (p< 0.05) by one-way ANOVA with Tukey’s post-hoc test.

### The accumulation of cysteine adducts responds to the changes in their precursors in central metabolism

Previous work from our group and others have shown that macrophages undergo highly dynamic changes upon classical activation. Particularly, glycolysis is strongly upregulated, and TCA cycle undergoes a two-stage remodeling driving by the sequential induction or inhibition of different enzymes in the TCA cycle^6,7^. This causes the dynamic changes of important intermediates derived from glycolysis and TCA cycle, including those that can give rise to the cysteine adducts (Fig. 5A).

Over the time-course of classical activation, we found that correlating with the increase in glycolytic intermediates like DHAP, all the glycolysis-derived cysteine adducts continue to increase, with the most abundant products (two isomers of C_6_H_12_NO_7_PS) accumulate to over 100-fold (Fig. 5B). In the TCA cycle, αKG level first decrease (due to early IDH inhibition^20^, then increase after 48 hours (due to strong OGDC inhibition at this time^6,7^). Showing strong correlation, we found αKG-Cys first decreases, then starts to accumulate significantly after 24-48 hours (Fig. 5C). Similar correlation was observed in Ita-Cys level. Ita-Cys first increase, then decrease (Fig. 5C), following the rise-and-fall of itaconate level (which is the result of early *Irg1* induction followed by later PDHC inhibition, as elucidated by previous research^6,7^). These results suggest the levels of αKG-Cys and Ita-Cys dynamically respond to the multi-stage remodeling of TCA cycle over the course of immune response. This is further confirmed by their pseudo-steady state labeling patterns. Over the activation time course, αKG-Cys labeling from glucose gradually decreases, following the labeling patten of αKG (Supp. Fig. 9A), consistent with the sequential IDH inhibition followed by OGDC inhibition driving the changes in αKG abundance, and subsequently αKG-Cys level. Ita-Cyc labeling largely tracks the labeling of itaconate, which decreases notably after 48h (Supp. Fig. 9B), when PDHC was found to be strongly inhibited^6^, restricting influx into itaconate production and downstream ITA-Cyc production.

To further validate that these newly identified cysteine adducts are produced in cells downstream of specific intermediates in glycolysis and TCA cycle and dynamically respond to changes in central metabolism, we examined a series of metabolic perturbations targeting central metabolism. In *Irg1* knock out cells, the production of itaconate is ablated. Consequently, the accumulation of Ita-Cys is ablated too (Fig. 5D). On the flip side, supplementing cells with itaconate resulted in increased Ita-Cys (Fig. 5E). The increase of glycolytic intermediates and the stronger accumulation of their cysteine adduct reflects the classical activation-induced increase in glycolytic rate. To perturb glycolytic rate, we either cultured cells in low glucose (1mM) environment, or treated cells with glycolysis inhibitor, 2-deoxyglucose (2-DG). Both treatments led to significant reduction of C_6_H_12_NO_7_PS isomers in activated macrophages, which is consistent with, but more sensitive than, the reduction in DHAP level (Fig. 5F-G). On the other hand, overexpressing glucose transporter GLUT1 in RAW264.7 macrophages increases the glucose uptake^21^. As a result, the accumulation of major glycolysis-derived cysteine-adducts is significantly increased, and the response is even more sensitive than the change in corresponding glycolytic intermediate (Fig. 5H-I).

### Increased cystine uptake supports the production of cysteine-adducts in stimulated macrophages

Cysteine is the common precursor of these classical activation-induced metabolites. Very intriguingly, we found that among all twenty amino acids, cysteine is uniquely increased in macrophages over the time course of classical activation (Fig. 6A), suggesting the possibility that a unique function of cysteine (such as supporting the production of cysteine adducts) rather than a shared function with other amino acids (such as synthesis of proteins and functional peptides) is particularly important in stimulated macrophages. Macrophages primarily obtain cysteine by taking up cystine from the media, through the transporter SLC7A11. We found *Slc7a11* is transcriptionally upregulated upon activation (Fig. 6B). *Slc7a11* is known to be transcriptionally regulated by NRF2^28,29^. We found that the expression of classical NRF2 targets all increases upon stimulation (Supp. Fig. 10), indicating NRF2 activation.

To test how cystine availability affects the production of cysteine adducts, we cultured BMDMs in RPMI media containing normal level of cystine (200 μM), high cystine (800 μM), or subjected to 48 hours of cystine depletion (0 μM). The substantial accumulation of all cysteine-adducts upon LPS/IFN stimulation, as observed in normal cystine condition, was eliminated in cystine depleted condition. On the flip side, high environmental cystine amplified the accumulation of these cysteine adducts in stimulated macrophages (Fig. 6C-I), showing a wide dynamic response range, while effect is much more profound for cystine depletion compared to high cystine perturbation. Normal cystine level in human circulation is around 30 μM (HMDB), much lower than standard RPMI media level. It is likely fluctuation of cystine level in physiological range has more profound effects.

### The production of cysteine adducts is regulated by NO

A characteristic metabolic function in classical activated macrophages is the production of nitric oxide (NO) via inducible nitric oxide synthase (iNOS). NO not only acts as an important pathogen killing agent but also plays significant regulatory roles in macrophages. Previous work from our group and others has demonstrated that NO is a regulator of glycolysis and TCA cycle^6,7,30–34^, which are upstream of cysteine adduct production. Furthermore, the production of NO and related reactive nitrogen species causes significant shifts in cellular redox environment in activated macrophages, which can influence the reaction between central metabolites and cysteine. Here we examined how NO impact the formation of these adducts by comparing the wildtype or iNOS knockout BMDMs with or without 48-hour classical activation.

Genetic knockout of iNOS, which eliminates the activation-induced NO production, significantly reduced the accumulation of most cysteine adducts derived from glycolysis upon stimulation (Fig. 7A). Conversely, treating iNOS knockout BMDMs with exogenous NO donor, DETA-NONOate, increased the stimulation-induced accumulation of most of these adduct (Fig. 7A). These results demonstrate NO is a positive regulator of glycolysis-derived cysteine adducts. The mechanism for this effect can be multifold. On one hand, NO is known to inhibit GADPH^35^, which enhances the accumulation of reactive glycolytic intermediates like DHAP upon stimulation. On the other hand, we found knockout of iNOS greatly reduced the upregulation of SLC7A11 upon classical activation (Fig. 7B), indicating NO also promotes the upregulation of cysteine supply upon stimulation. To note, however, NO donor treatment itself is not sufficient to cause large accumulation of these cysteine adduct, immune stimulation with LPS and IFNγ is required (Fig. 7A). This indicates NO is a regulator tuning this stimulation-induced metabolic response, but it is not the sole driver.

In contrast to the glycolysis-derived adducts, the stimulation-induced accumulation of Ita-Cys is further enhanced by iNOS knockout (Fig. 7C). Our previous work had shown that NO can drive the decrease of itaconate at the later part of classical activation time course by inhibiting PDHC^6^. Consistently, the increased accumulation of Ita-Cys in iNOS knockout cells at this time point (48-hours post stimulation) mirrors the persistent accumulation of itaconate in iNOS KO cells (Fig. 7D).

In sum, results above showed that NO is a regulator tuning the accumulation of cysteine adducts. This can be a combined effect of NO regulating glycolysis and TCA cycle, regulating cystine uptake, and leading to the increase of reactive oxygen and nitrogen species (RONS), which are free radicals known to impact the reactivity of thiols.

### The newly identified cysteine-adducts are present in human samples and increase in inflammatory conditions

Finally, we seek to validate whether these metabolites discovered in cultured murine macrophages are present in human tissues and primary human cells. We performed LCMS-based metabolomic analysis of granuloma annulare (GA), an inflammatory skin condition characterized by the formation of granulomas in the skin^36^. Granulomas are comprised of aggregating macrophages and other immune cells ^37,38^. Pairs of skin specimens were collected from GA lesion and non-lesional skin (termed ‘healthy’) contralateral to the GA skin biopsy from the same patient (Fig. 8A). Most of the newly discovered cysteine adducts were found in the patient skin samples and are significantly increased in GA lesion compared to contralateral healthy controls (Fig. 8B-E). The exceptions are those two resulted from addition reaction: Ita-Cys was not reliably detected in these samples and Suc-Cys does not change significantly (Fig. 8F). These results confirmed that the cysteine-adducts, especially those novel metabolites resulted from reactions with carbonyl-containing intermediates in central metabolism, are human metabolites and are associated with inflammatory condition.

**Figure 8.**
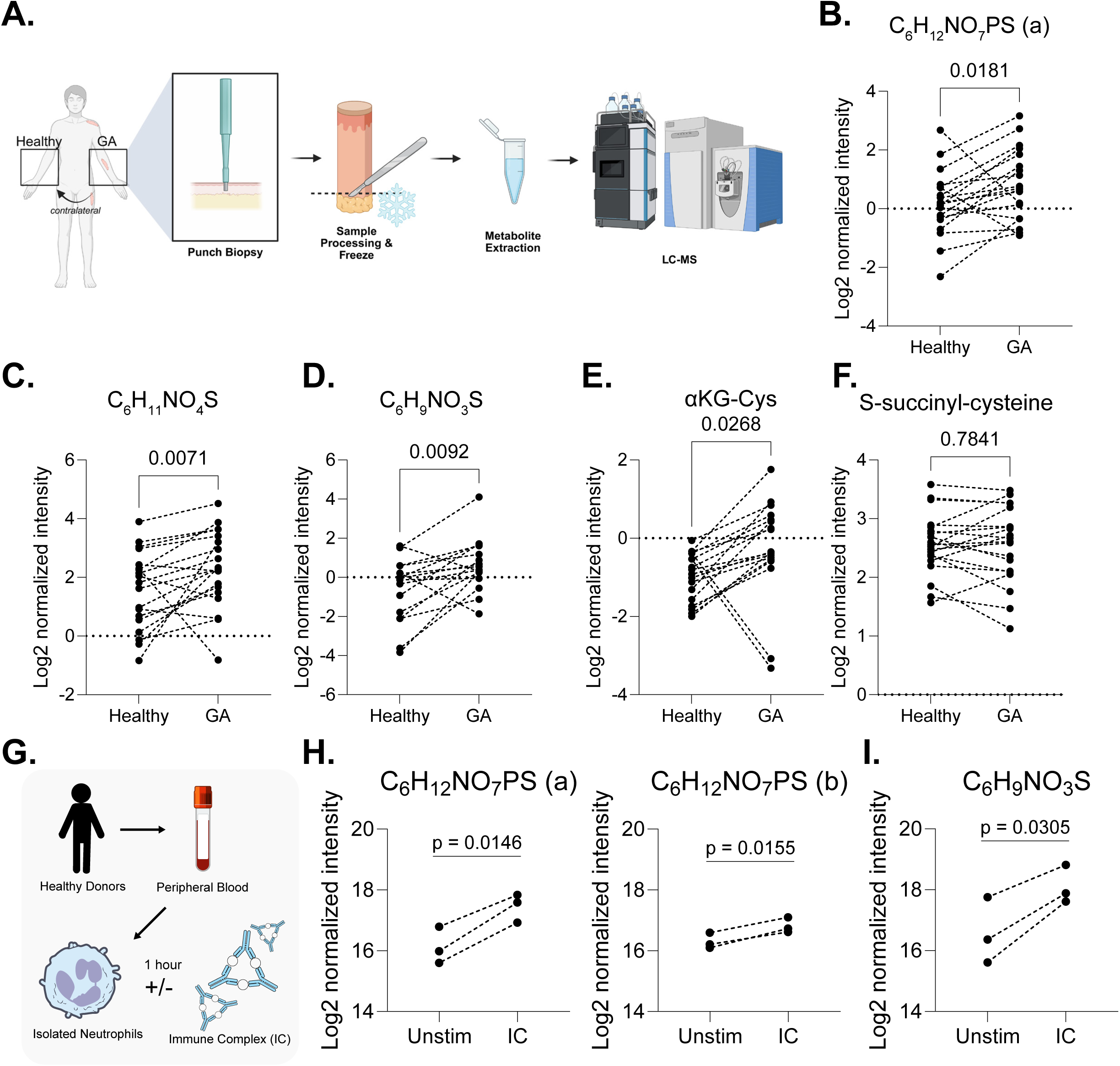
Cysteine adducts are elevated in inflammatory skin disease and activated human neutrophils. **A.** Schematic of granuloma annulare (GA) and paired non-lesional (‘healthy’) skin sample collection and processing. Samples were collected from 20 patients. **B-F.** Log2 normalized intensities of major central carbon metabolite-cysteine adducts in paired healthy and GA skin samples. Statistical comparisons were performed using Wilcoxon matched-pairs signed rank test with exact p-value reported. **G.** Schematic of human peripheral blood neutrophils stimulated with immune complexes (IC). Primary peripheral blood neutrophils were freshly isolated from 3 healthy donors **H-I.** Elevation of major glycolysis-derived cysteine-adducts in human peripheral blood neutrophils upon stimulation with ICs for 1h. Mean +/- SD, n = 3. Log2 normalized intensities of major central carbon metabolite-cysteine adducts in paired unstimulated and IC-stimulated neutrophils from the same donor. Statistical comparisons were performed using paired two-tailed t-tests.

Moreover, we found the production of these metabolites is not limited to macrophages. In human peripheral blood neutrophils, activation by immune complexes (IC) caused both C_6_H_12_NO_7_PS isomers and C_6_H_9_NO_3_S (all major cysteine-adducts derived from glycolysis) to increase significantly within only 1 hour (Fig. 8G-I), suggesting their accumulation may be more broadly associated with innate immune cells activation.

## DISCUSSION

In this work, we identified metabolites that are produced from reactions between free cysteine and reactive intermediates in central carbon metabolism, including glycolysis intermediates (i) GAP, (ii) DHAP, (iii) MGO, and TCA cycle-derived metabolites, (iv) itaconate, (v) αKG, and (vi) fumarate. Most of the dozen of products have never been reported in mamlian metabolome before, the reactions of GAP, DHAP and αKG with cysteine are especially novel. These central metabolite-cysteine adducts accumulate substantially in macrophage upon inflammatory activation. An open question is why these metabolites specifically accumulate in such condition.

Interestingly, we noticed most of the precursors forming cysteine-adducts have been found to directly cause protein post-translational modifications (PTM)^39^, many of which are formed non-enzymatically. Some of these PTMs can be damaging to cells, while many also have important regulatory roles^10,26,40–42^. The close chemical relationship between these important PTMs and cysteine adducts leads to the hypothesis that one significance for the formation of cysteine adducts may be to buffer non-enzymatic protein PTMs. Moreover, some of the cysteine adducts may even be formed through reactions where free cysteine mediates the removal of corresponding PTMs on protein, performing a similar function better known to be carried by glutathione. Indeed, an inspiring recent work has demonstrated an example that cysteine has Glo1 and Glo2 activity towards MGO-derived modification^22^. During immune activation, metabolic demands greatly increases and cells switch to a much more reactive metabolic state. For instance, glycolysis is upregulated and reactive glycolytic intermediates with protein modifying potential accumulates. The buffering could be particularly important in this situation.

Consistent with the idea that increased cysteine adduct production is likely an active response upon classical activation, we found macrophages uniquely upregulated the import of cysteine among all amino acids. Chemically, cysteine is unique among all amino acids in that it contains a reactive thiol. While compared to other thiol-containing low molecular weight compounds (LMW-SH) like glutathione, or cysteine residues in proteins, free cysteine also contains an amine group that is three bonds away. The combination of thiol and amine in the same molecule allows intermolecular S- to N-transfer and formation of intermolecular ring, generating more stable structures that do not further transfer modifications (unlike S-lactoyl-glutathione can transfer lactoyl-modification to lysine residue of proteins^42^), potentially making cysteine a unique buffer compound to sequester reactive molecules. Indeed, our solved cyclic structure of αKG-Cys support the direct participation of both thiol and amine in the reaction. Furthermore, based on the most abundant and fast-producing products between cysteine and DHAP, GAP, or αKG (C_6_H_12_NO_7_PS, and C_8_H_11_NO_6_S), we searched whether similar dehydration condensation products could be produced between these reactive central metabolites and glutathione, but did not detect any corresponding product. Biologically, cysteine also stands out among all LMW-SH for its high availability. All these chemical and biological properties make cysteine a unique reactant in cells and can potentially play a key role in regulating the non-enzymatic protein PTMs by reactive metabolites. Interestingly, the protein best known for being regulated by non-enzymatic PTM driven by these metabolites is KEAP1, which is the suppressor of NRF2 and an important regulator in macrophage immune response. Both GAP and MGO can cause S-lactoylation on cysteine residue of KEAP1^41^, additionally, it was found that MGO can cause a methylimidazole crosslink between proximal cysteine and arginine residues (MICA) in the KEAP1 protein. Itaconate and fumarate can cause S-itaconylation and S-succinylation of KEAP1, respectively^10,27^. All of these modifications ultimately lead to the release and activation of NRF2. Interestingly, we observed that upon LPS/IFN stimulation, the NRF2 targets broadly increased, consistent with previous reports and the stimulation-induced accumulation of KEAP1-modifying metabolites. Limiting extracellular cystine further exacerbated the increase of NRF2 activity (Supp. Fig. 10), consistent with (though does not provide direct evidence) that cysteine could provide a buffer for non-enzymatic regulatory KEAP1 PTMs, tempering NRF2 activation. Intriguingly, NRF2 is also the upstream regulator of *Slc7a11,* and we observed *Slc7a11* showed a strong increase upon cystine depletion (Supp. Fig. 10). The regulation of *Slc7a11* by NRF2 could potentially provide a negative feedback regulation loop, where KEAP1 senses the accumulation of reactive metabolites upon classical activation, and in turn upregulate *Slc7a11* to increase cysteine supply to counter the reactive stress and lower PTMs by these reactive metabolites, keeping cellular homeostasis and preventing NRF2 hyper activation. This hypothesis merits future work for in-depth validation.

Besides keeping dynamic homeostasis, it is also possible the adducts of central carbon metabolites and cysteine have signaling/regulatory effects or anti-microbial effect themselves. Some other central metabolites-amino acid adducts have been recently found to have important regulatory roles. A particularly intriguing example is N-lactoyl-phenylalanine (Lac-Phe), a signaling metabolite from lactate. It was suggested that by conjugating a glycolytic metabolite reflecting metabolic state and nutrient availability (lactate) to an amino acid to form a signaling molecule lasting longer than lactate itself, it generates a more dynamically suitable signal to regulate feeding and downstream physiological responses^43^. Similarly, the cysteine adducts we elucidate here has similar properties, they closely reflect the respective changes in central metabolism (specifically the upregulation of glycolysis and the dynamic reprogramming of TCA cycle associated with macrophage activation), while the increase is of greater amplitude and dynamically the signal of some adducts can last longer than their precursors, as exemplified by Ita-Cys continuing to be elevated after itaconate itself fall back down. This can make them sensitive and relatively stable markers to reflect that the prior macrophage activation and metabolic reprogramming, in a way, reflect a short metabolic memory. The potential signaling/regulatory role of these compounds is of interest for further investigation.

This work also points to intriguing questions about the metabolism of these newly identified central metabolite-cysteine adducts. Regarding their production, while here we showed these reactions *can* occur without an enzyme, underscoring the high chemical and thermodynamic feasibility, it is also possible enzymes can catalyze some of the reactions, especially given our kinetic tracing experiment showed that many of these compounds (particularly those derived from glycolysis) turnover fast, with half-life ranging from less than 30 minutes to a couple of hours. Special biochemical environment, such as low pH or high RONS in certain compartments in activated macrophages, can also accelerate these reactions. Furthermore, despite their rapid turnover rates, we did not detect significant excretion of these compounds in spend media, suggesting there likely are active mechanisms to catabolize these compounds. These questions merit further investigation.

From a methodological perspective, this work exemplifies a framework to identify interesting, previously unknown metabolites and biochemically map them into the uncharted part of metabolic network. The framework starts with discovery of mass spectrometry (independent of metabolite matching to existing metabolite database), integrating the use of exact mass, natural isotopic distribution, MS/MS, and isotopic tracing to identify the formula and sources of compounds with interesting biological characteristics (e.g., uniquely associated with the classical activation state in macrophages, or change dynamically upon stimulation). Then in vitro investigation, NMR analysis, and correlation analyses within the metabolomic datasets across varieties of metabolic perturbation conditions (e.g., inhibition or knockdown of metabolic enzymes, transporter, perturbations by NO) were applied to rigorously verify the reaction making these compounds and their connections with known parts of metabolic network. This workflow can be applied to similar questions beyond macrophage activation, such as identifying metabolites uniquely produced in a specific genetic condition or mapping out previously unknown reactions that are greatly increased in other specific cellular state. This is valuable in identifying biomarkers and providing insights into how cells handle the metabolic need and challenge associated with a specific environment or functional state.

## METHODS

### Primary Bone Marrow Derived Macrophages

As previously described^6,30,33,44^, primary bone marrow-derived macrophages (BMDMs) were isolated from 8–12-week-old male and female wild-type C57BL/6J mice or B6.129P2-Nos2^tm1Lau^/J (iNOS KO) mice or homozygous C57BL/6NJ-Acod1*^em^*^1^*^(IMPC)J^*/J *(Irg1* KO) mice (Jackson Laboratory). All animal procedures were approved by the University of Wisconsin-Madison Institutional Animal Care and Use Committee. Mice were group-housed on a 12-hour light/dark cycle at 18-23C with 40-60% humidity and provided food and water ad libitum. Bone marrow was flushed from femurs and tibias and pooled. Cells were differentiated into macrophages on petri dishes in RPMI 1640 medium containing L-glutamine (VWR, 16750-070), 10% FBS (Hyclone, SH30910.03), 25 mM HEPES, 1% penicillin/streptomycin (Gibco, 15-140-122), and fibroblast-conditioned medium (as a source of M-CSF). Media was replaced on days 3, 5, and 6. On day 7, differentiated BMDMs were harvested, seeded, and cultured in RPMI 1640 complete medium supplemented with 20 ng/mL recombinant mouse M-CSF (R&D Systems, 416-ML-050). For classical activation, BMDMs were treated with 50 ng/mL LPS (E. coli O111:B4; Sigma-Aldrich, L3024) and 30 ng/mL recombinant mouse IFNγ (R&D Systems, 485-MI-100) for the indicated durations. For alternative activation, BMDMs were treated with with 20 ng ml^−1^ IL-4 (cat. no. 404-ML, R&D Systems) and 20 ng ml^−1^ IL-13 (cat. no. 413-ML-005, R&D Systems).

For exogenous NO treatment, 200 µM diethylenetriamine NONOate (DETA-NONOate; Cayman Chemical, 82120) was added and maintained for the indicated durations. Primary cell cultures were maintained at 37°C in a humidified atmosphere with 5% CO_₂_.

For L-cystine depletion experiments, RPMI-1604 without L-cystine, L-glutamine, and L-methionine (Millipore, R7513-100ML) was used and then supplemented with formulation concentrations of glutamine (Fisher Scientific, BP379-100) and methionine (Sigma, M5308-25G); cystine (Millipore, C7602-25G) was supplemented at the reported experimental concentrations or kept absent.

### RAW264.7 Cell Culture

GLUT1 overexpression RAW264.7 cells (Gift from Lisa Makowski, PhD at UNC-Chapel Hill; Empty Vector clone, rat GLUT1 OE clone as reported in Freemerman et al.^21^) were cultured in DMEM with high glucose with 10% FBS and 1% penicillin/streptomycin and maintained under blasticidin selection conditions (10 µg/mL). For glycolysis inhibition, 2-deoxyglucose (1 mM) was supplemented. For low glucose conditions, glucose-free media was supplemented with 1 mM glucose. Cell cultures were maintained at 37°C in a humidified atmosphere with 5% CO_₂_. For classical activation, RAW264.7 cells were treated with 50 ng/mL LPS (E. coli O111:B4; Sigma-Aldrich, L3024) and 10 ng/mL recombinant mouse IFNγ (R&D Systems, 485-MI-100)

### Stable Isotope Tracing

For stable isotope experiments, the following isotopes were used: 3-3-^13^C_2_-L-cystine (Cambridge Isotope Laboratories, CLM-1822-H), U-^13^C-D-glucose (Cambridge Isotope Laboratories, CLM-1369-1), U-^13^C-L-glutamine (Cambridge Isotope Laboratories, CLM-520-0). Each respective isotope was used at RPMI-1640 formulation concentrations in cystine-, glucose-, or glutamine-deplete media, respectively.

For the symmetric isotope labeling experiment, BMDMs were seeded and cultured in full RPMI-1640 described above. The cells were stimulated with LPS and IFNγ. After 24-hours, the media was replaced with full unlabeled RPMI media or full RPMI-1640 media containing isotope tracer, with the stimulus at the same concentration. After additional 24-hours, the intracellular metabolites were extracted from the cells. For the kinetic labeling, BMDMs were seeded and cultured in full unlabeled culture media described above. The cells were stimulated with LPS and IFNγ. After 48-hours stimulation, media was switched to chemically identical media but containing U-13C-glucose instead of ^12^C-glucose, for the indicated duration of time, then the intracellular metabolites were extracted.

### In vitro reactions

The following substrates were used for in vitro reactions: L-cysteine (Sigma Aldrich, 30089), alpha-ketoglutaric acid disodium salt dyhydrate (Sigma Aldrich, 75892-25G), itaconic acid (Millipore, I29204-100G), D-glyceraldheyde-3-phosphate (MedChemExpress, HY-151223), dihydroxyacetone phosphate lithium (Millipore, 37442-100MG-F), methylglyoxal solution (Millipore, M0252).

All substrates were prepared in reaction buffer (0.2 mM NaHCO₃, pH 7.4). Substrates were mixed to the final concentrations reported in each experiment and incubated at room temperature or 37°C as indicated. Reaction aliquots at designated timepoints (1 and 24 hours) were diluted 1:20 (v/v) in LC-MS-grade H₂O prior to LC-MS analysis.

### Metabolite extraction and LC-MS analysis

To measure intracellular metabolites, cells were washed twice with D-PBS and metabolites were extracted with cold LC-MS grade 80:20 (v/v) methanol:water (Thermo Scientific, A4564, W64). The resulting pellet was extracted twice and the extracts were pooled in a tube and dried under a nitrogen stream. Dried samples were resuspended in LC-MS grade H_2_O. Samples were analyzed using a Q Exactive Orbitrap Mass Spectrometer (Thermo Fisher Scientific) coupled to a Vanquish Horizon UHPLC System. Xcalibur 4.0 was used for data acquisition. Samples were separated on a 100 × 2.1 mm, 1.7 μM ACQUITY UPLC BEH C18 Column (Waters) with a gradient of solvent A (97:3 (v/v) water:methanol, 10LmM TBA (cat. no. 90781, Sigma-Aldrich)), 9 mM acetic acid (pH 8.2, cat. no. A11350, Thermo Fisher Scientific) and solvent B (100% methanol). The gradient was: 0 min, 5% B; 2.5 min, 5% B; 17 min, 95% B; 21 min, 95% B; 21.5 min, 5% B. The flow rate was 0.2 ml min^−1^ with a 30 °C column temperature. Data were collected in full scan negative mode at a resolution of 70 K. Settings for the ion source were: 10 auxiliary gas flow rate, 35 sheath gas flow rate, 2 sweep gas flow rate, 3.2 kV spray voltage, 320 °C capillary temperature and 300°C heater temperature. For MS/MS analysis, the parameters were: isolation window 0.7, step collision energy of 20, 30, 40 NCE.

Data were analyzed with MAVEN (v.2011.6.17,for MS1 level quantification) and Mzmine (V. 7.8.30, mzio GmbH). Metabolite levels were normalized to total protein content quantified using Pierce™ BCA Protein Assay Kit (Thermo Scientific, 23227) or by cell area using an IncuCyte S3 Live-Cell Analysis System (Sartorius). For targeted analysis and analysis of known metabolites (e.g., amino acids in Fig. 6), metabolites were identified based on exact *m/z* and retention times determined using standards. For untargeted analysis (Figure 1), LCMS features were filtered to retain those present in at least 50% of samples within at least one condition. Missing values were imputed at one-fifth of the minimum detected value. Mean intensities were calculated for each condition, and fold changes were determined. Statistical significance was assessed using unpaired two-tailed t-tests. Features showing >2.5-fold increase in stimulated vs unstimulated conditions with p < 0.05 were classified as activation-specific.

¹³C isotopes were identified as features with m/z + 1.0034 ± 0.015 Da and retention time ± 0.25 min of parent peaks, with relative intensity 0.5-20%. Possible sulfur-containing metabolites were identified by ³⁴S isotopes (m/z + 1.9958 ± 0.015 Da, RT ± 0.25 min) with characteristic 2-6% natural abundance relative to parent peaks. For cystine labeling analysis, ¹²C-¹³C pairs were identified based on the expected mass shift (1.0034 Da), reciprocal ratio changes (unlabeled/labeled >1.2 for ¹²C parent, <0.8 for ¹³C isotope), and p < 0.05.

### NMR Data Acquisition

All NMR experiments were performed on a 500 MHz spectrometer equipped with a TCO cryoprobe. ^1^H NMR spectra were collected using the zgesgp water-suppression pulse sequence. Free induction decays (FIDs) were acquired with 32,000 complex points and 256 scans, using a 16 ppm spectral width centered at 4.7 ppm. ^1^H-^1^H COSY was acquired with 32 scans per increment and 512 increments in the indirect dimension. The spectral width was 10 ppm, centered at 4.7 ppm. ^13^C NMR spectra were collected with 2,048 scans over a 200 ppm spectral width centered at 100 ppm. ^13^C DEPT-135 spectra were acquired with 1,024 scans and a 160 ppm spectral width centered at 80 ppm. ^1^H-^13^C HSQC spectra were acquired with 32 scans, 512 increments, and a 10 ppm spectral width centered at 4.7 ppm for the ^1^H dimension. The ^13^C spectral width was 200 ppm, centered at 100 ppm.

### Isolation, culture and stimulation of human peripheral blood neutrophils

Human neutrophils were isolated from peripheral blood freshly collected from healthy donors, following the protocol approved by the University of Wisconsin Institutional Review Board (protocol no. 2019-1031). Informed consent was obtained from all participants. Neutrophils were purified using the MACSxpress Whole Blood Neutrophil Isolation Kit (Miltenyi Biotec, 130-104-434) followed by erythrocyte depletion (Miltenyi Biotec, 130-098-196) according to the manufacturer’s instructions. Neutrophils were spun down at 300 x *g* for 5 minutes, resuspended in neutrophil culture media – RPMI 1640 w/o glucose (ThermoFisher, 11879020) supplemented with 5 mM glucose (Fisher Scientific, D16500) with 10% human plasma – and kept at 37°C in incubators under 5% CO_2_. Purity and cell viability of isolated cells has been previously verified^45,46^. The human neutrophils were stimulated for one hour with 1 μl of OVA-ICs per 50,000 cells. OVA-ICs were generated as previously described^47^. To extract intracellular metabolites, cells were extracted with 80 μls of cold LC-MS-grade acetonitrile/methanol/water (40:40:20 v:v:v). After the extraction, samples were dried under a stream of nitrogen gas for LCMS analysis.

### Human skin tissue sample collection and processing for metabolomics

Skin specimens were obtained under local anesthesia (lidocaine) with a 6 mm disposable biopsy punch. For each subject, 2 specimens were collected. One from an active GA lesion and one from healthy skin contralateral to the GA skin biopsy. All human studies were approved by the authors’ institutional review board at the University of Wisconsin-Madison (protocol no. 2021-0047), and patients gave their written, informed consent. The study procedures, including obtaining written informed consent, were performed in accordance with the ethical standards of the Helsinki Declaration of 1975.

Skin biopsies were collected from 20 subjects with biopsy confirmed granuloma annulare after local anesthesia. Samples were washed in a petri dish containing phosphate buffered saline to clean off debris and blood. Subcutaneous fat was removed with a scalpel. Samples were dried using kimwipes and immediately flash frozen in liquid nitrogen. Samples were then pulverized using a biopulverizer cooled with dry ice. Powdered sample was transferred to a 2mL tube and 400µl of chilled 3:1 butanol:methanol with 10µM internal standards (^3^D-2-hydroxyglutarate and ^15^N4-arginine) was added. Samples were vortexed on a tissue disruptor for 1 minute and then sonicated on ice for 5 s followed by 5 s rest for a total of two times at 60% amplitude with a probe sonicator. Then, 400µl of chilled 3:1 heptane:ethyl acetate was added, and samples were vortexed and sonicated again. Lastly, 400µl of chilled 1% acetic acid was added, and samples were vortexed and sonicated one last time. Samples were spun down and the aqueous phase was transferred to a new 1.5mL tube. Samples were dried under nitrogen flow and then resuspended in LCMS grade water relative to protein content of the biopsy (100µl LCMS water/40µg protein). LCMS analysis and downstream data analysis was performed using similar approach as described above. Low detection sample pairs were filtered out if intensity of a compound was low (<3e4 ion count) in both healthy and GA samples of the same patient. Peak intensities were normalized by median peak intensity values and then log2-transformed.

### Quantitative PCR

Cells were washed with phosphate-buffered saline (Thermo Scientific, AAJ67802-K2) and lysed with RNA Stat60 (Fisher Scientific, NC9489785) with subsequent RNA isolation performed per manufacture’s protocol. Chloroform (Sigma Aldrich, 650498) was used for phase separation. RNA was precipitated with isopropanol (Thermo Scientific, AC327272500) and washed with 75% [v/v] molecular-grade ethanol (VWR, 71006-012). RNA pellets were resuspended in molecular-grade water and quantified with a NanoDrop 2000c Spectrophotomer (Thermo Scientific). cDNA was synthesized using the High-Capacity cDNA Reverse Transcription Kit (Thermo Scientific, 4368813). Quantitative PCR reactions were performed with 10 ng of cDNA per reaction using PowerUp™ SYBR™ Green Master Mix (Thermo Scientific, A25777) on an Applied Biosystems QuantStudio 7 Pro Real-Time PCR System (Thermo Scientific). Gene expression was analyzed using the ΔΔCt method with *Hnrpab* as the reference gene. The following target primers were used: *Slc7a11* - Forward: 5’-GGCACCGTCATCGGATCAG-3’, Reverse: 5’-CTCCACAGGCAGACCAGAAAA-3’; *Nqo1* - Forward: 5’- AGGATGGGAGGTACTCGAATC-3’; Reverse: 5’- AGGCGTCCTTCCTTATATGCTA-3’; *Hmox1*, Forward: 5’-AAGCCGAGAATGCTGAGTTCA-3’, Reverse: 5’-GCCGTGTAGATATGGTACAAGGA-3’; *Gclm* - Forward: 5’- AGGAGCTTCGGGACTGTATCC-3’; Reverse: 5’- GGGACATGGTGCATTCCAAAA-3’; *Hnprab* - Forward: 5’- AGGACGCGGGAAAAATGTTC-3’; Reverse: 5’-CAGTCAACAACCTCTCCAAACT-3’.

### Software and Statistical Analysis

LC-MS data analysis was completed with Maven (v6.2). Chromatograms were generated using mzmine (v4.8.30, mzio GmbH). Statistical analyses were completed in GraphPad Prism (v10) for Windows (Graph Pad Software) or were performed in R (version 4.5.1.) using tidyverse (v2.0.0.), ggplot2 (v4.0.2.), and ggrepel (v0.9.6.) packages. Chemical structures were generated using ChemDraw Professional (v 22.0.0.22). Final figures were created with Adobe Illustrator 2025. Figure 8A was created in BioRender. Arp, N. (2026) https://BioRender.com/69mzrg6. Methods for statistical analyses are noted in the corresponding figure legends.

## DATA AVAILABILTY

All data reported in this manuscript is available on request.

## ACKNOWLEDGEMENTS

The authors thank Lisa Makowski, PhD at the University of North Caroline-Chapel Hill for sharing GLUT1 overexpressing cell lines.

---

**Supplemental Figure 1.**
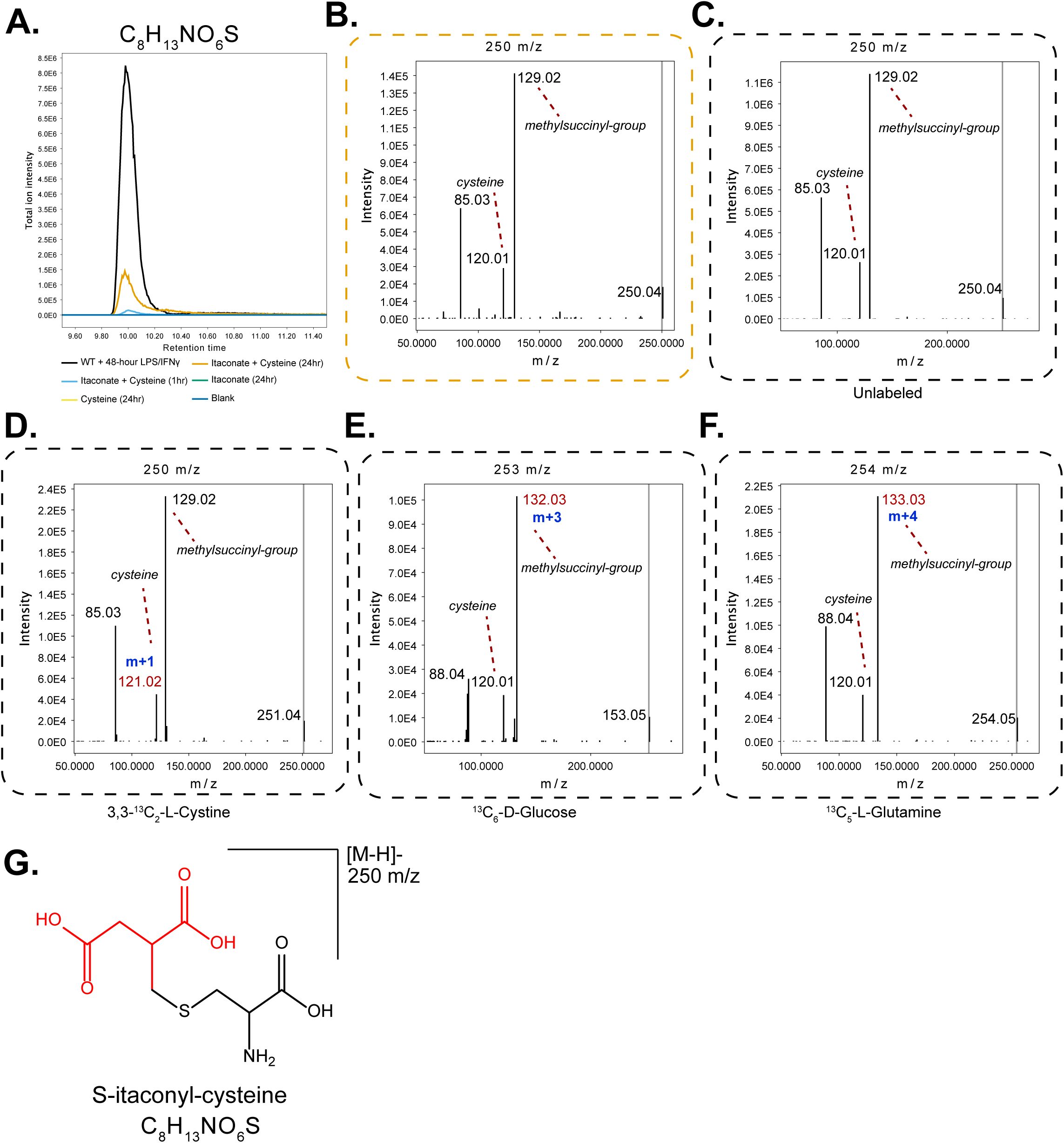
Chromatogram and MS/MS fragmentation patterns of S-itaconyl-cysteine (C_8_H_13_NO_6_S). **A.** EIC at 250.039 m/z showing S-itaconyl-cysteine formation from itaconate and cysteine coincubations (1 and 24 hours) compared that detected in LPS/IFNγ-stimulated BMDMs (black trace). **B-F.** MS/MS spectra correspond to the MS1 peak shown in A, in **B**) reaction mix of itaconate and cysteine in vitro coincubation (**C**) cell extract of unlabeled LPS/IFNγ-stimulated BMDMs, (**D**) cell extract of 3,3-^13^C_2_-L-cystine-labeled LPS/IFNγ-stimulated BMDMs, (**E**) cell extract of ^13^C_6_-D-glucose-labeled LPS/IFNγ-stimulated BMDMs, (**F**) cell extract of ^13^C_5_-L-glutamine-labeled LPS/IFNγ-stimulated BMDMs. The MS/MS show fragments correspond to itaconyl-group and cysteine with the expected mass shift from cystine, glucose, or glutamine tracer. **G.** Proposed structure of S-itaconyl-cysteine.

**Supplemental Figure 2.**
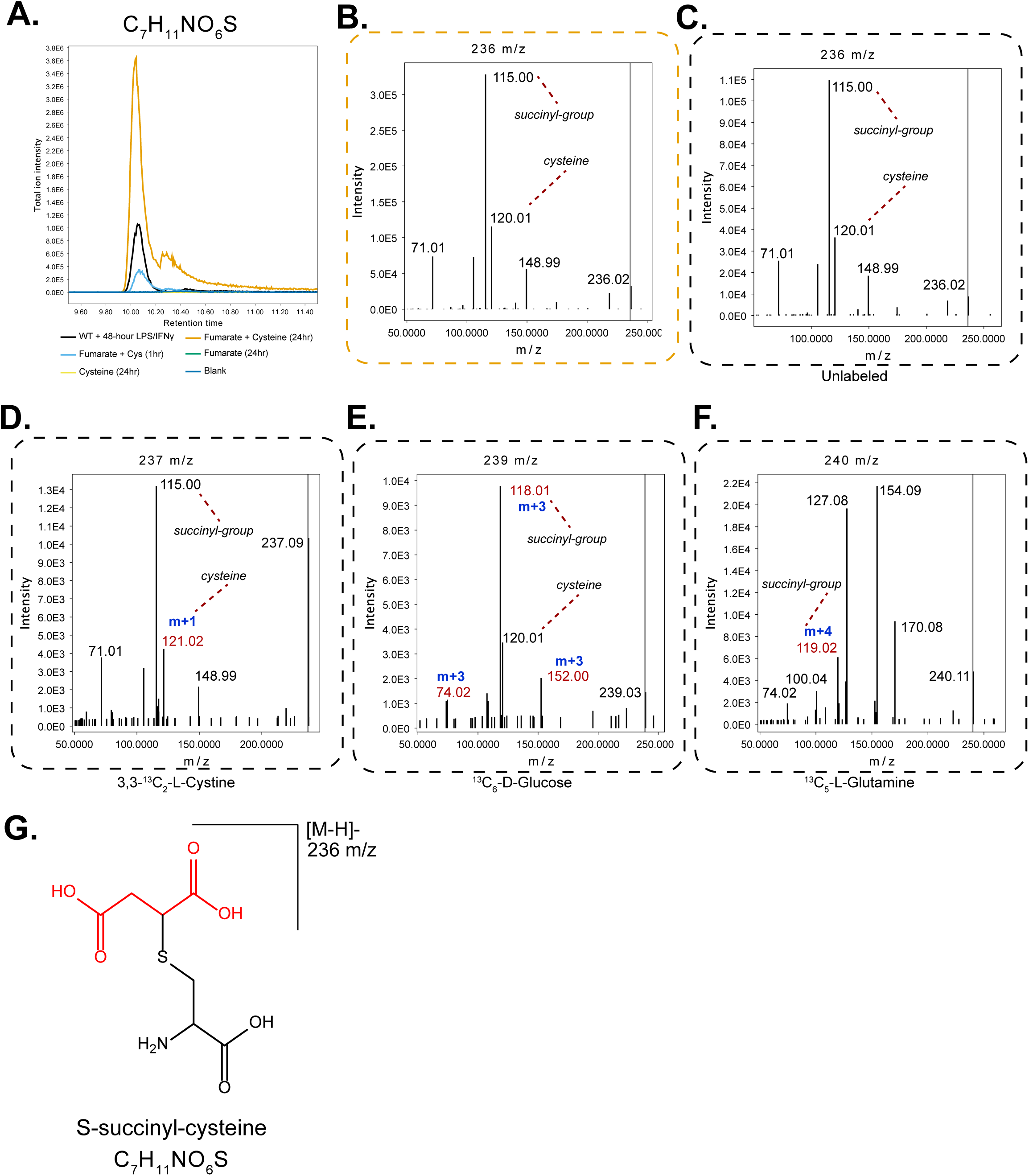
Chromatogram and MS/MS fragmentation patterns of S-succinyl-cysteine (C_7_H_11_NO_6_S). **A.** EIC at 236.023 m/z showing S-succinyl-cysteine formation from fumarate and cysteine in vitro coincubations (1 and 24 hours) compared to LPS/IFNγ-stimulated BMDMs (black trace). **B-F.** MS/MS spectra correspond to the MS1 peak shown in A, in **B**) reaction mix of fumarate and cysteine in vitro coincubation (**C**) cell extract of unlabeled LPS/IFNγ-stimulated BMDMs, (**D**) cell extract of 3,3-^13^C_2_-L-cystine-labeled LPS/IFNγ-stimulated BMDMs, (**E**) cell extract of ^13^C_6_-D-glucose-labeled LPS/IFNγ-stimulated BMDMs, (**F**) cell extract of ^13^C_5_-L-glutamine-labeled LPS/IFNγ-stimulated BMDMs. The MS/MS show fragments correspond to succinyl-group and cysteine with the expected mass shift from cystine, glucose, or glutamine tracer. **G.** Proposed structure of S-succinyl-cysteine.

**Supplemental Figure 3.**
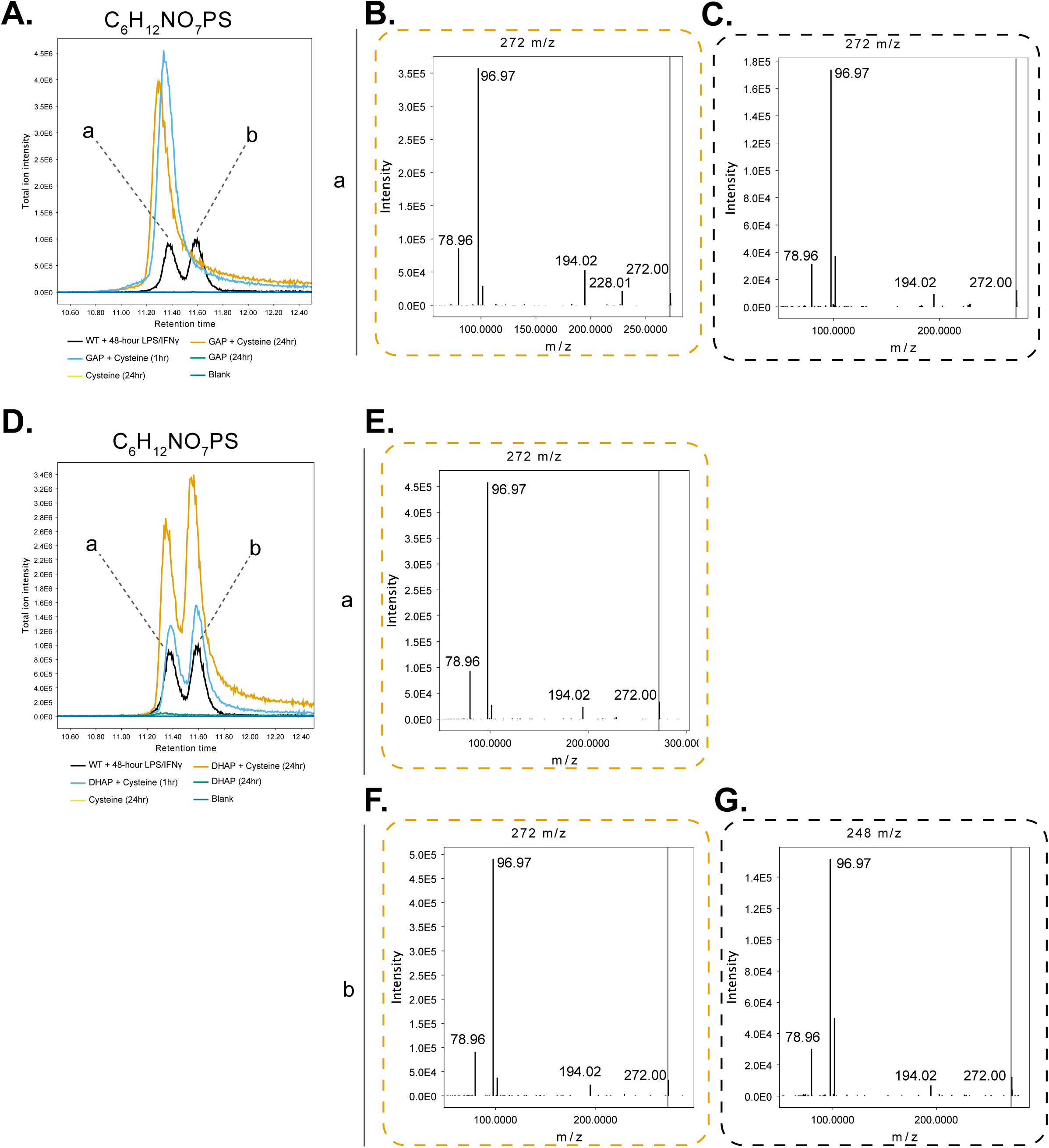
Chromatogram and MS/MS fragmentation patterns of C_6_H_12_NO_7_PS. **A.** EIC at 272.000 m/z showing C_6_H_12_NO_7_PS formation from: GAP + cysteine in vitro co-incubation (producing isomer a) compared in LPS/IFNγ-stimulated BMDMs (black trace, containing isomers a/b). **B-C.** MS/MS fragmentation spectra of parent mass 272 at RT=11.4 (isomer a) in reaction mix of GAP and cysteine in vitro coincubation (**B**) or cell extract of LPS/IFNγ-stimulated BMDMs (**C**). **D.** EIC at 272.000 m/z showing C_6_H_12_NO_7_PS formation from: DHAP + cysteine coincubation (producing isomers a/b) compared to LPS/IFNγ-stimulated BMDMs (black trace, containing isomers a/b). **E-F.** MS/MS fragmentation spectra of parent mass 272 at RT=11.4 (isomer a, E) or at RT=11.6 (isomer b, F) in reaction mix of DHAP + cysteine co-incubation. **G** MS/MS fragmentation spectra of parent mass 272 at RT=11.6 (isomer b) in cell extract of LPS/IFNγ-stimulated BMDMs.

**Supplemental Figure 4.**
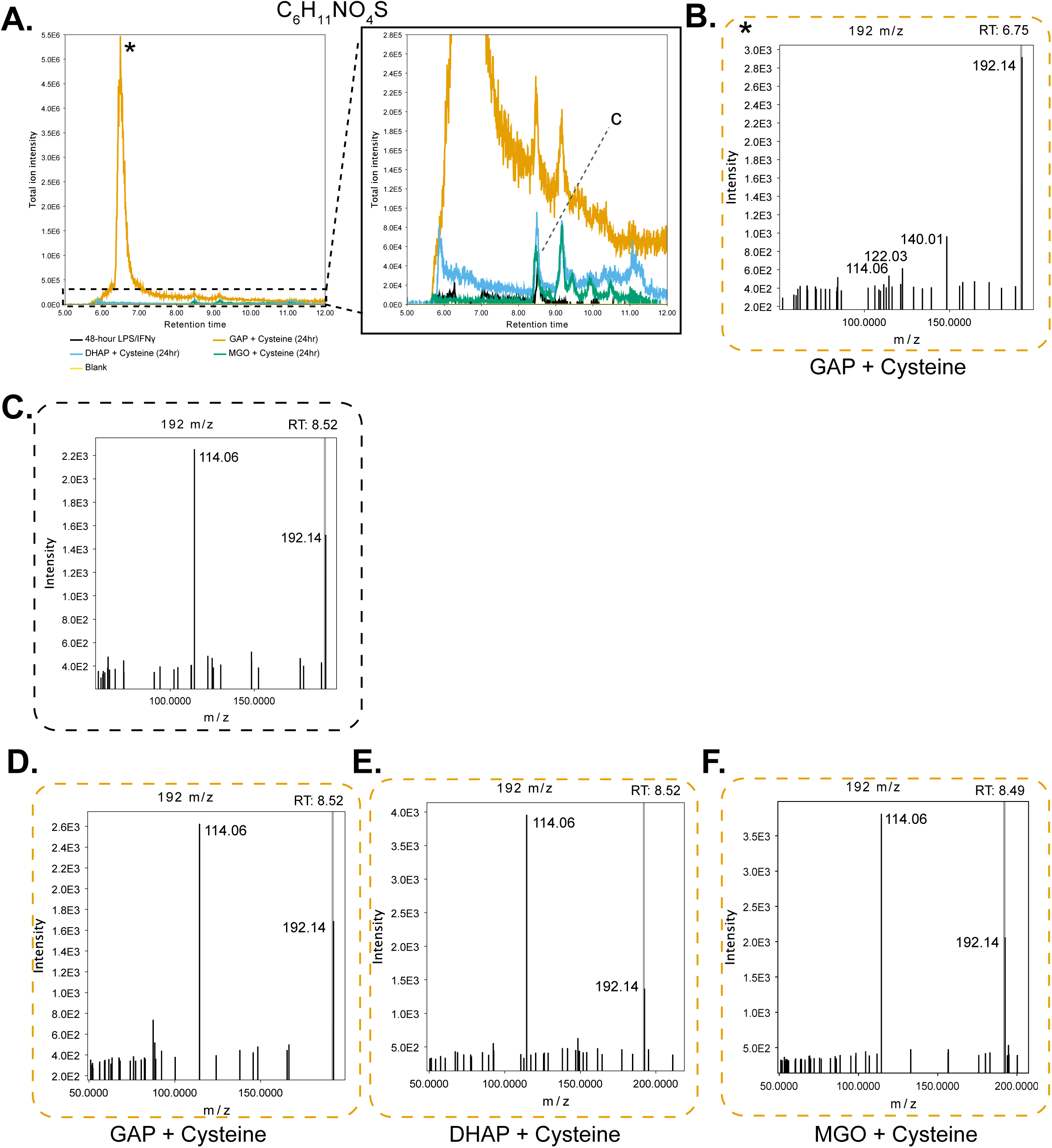
LC-MS/MS characterization of C_6_H_11_NO_4_S isomers. **A.** EIC at 192.034 m/z showing C_6_H_11_NO_4_S formation from DHAP (blue), GAP (yellow), or MGO (green) reacting with cysteine in vitro (24 hours), compared to LPS/IFNγ-stimulated BMDMs (black trace). As the product from GAP and cysteine co-incubation is very high in abundance compared to other conditions, the chromatogram is zoomed in on the right to better show all the isomers from all samples. **B** MS/MS spectra of parent mass 192, at RT=6.75 in the reaction mix of GAP and cysteine 24h in vitro co-incubation (the most abundant isomer from this reaction). **C-F.** MS/MS spectra of parent mass 192, at RT=8.5 in (**C**) cell extract of LPS/IFNγ-stimulated BMDMs, (**D**) GAP and cysteine in vitro co-incubation, (**E**) DHAP and cysteine in vitro co-incubation, (**F**) MGO and cysteine in vitro co-incubation. This peak corresponds to the most abundant isomer observed in cells (as indicated as isomer c in Figure 1).

**Supplemental Figure 5.**
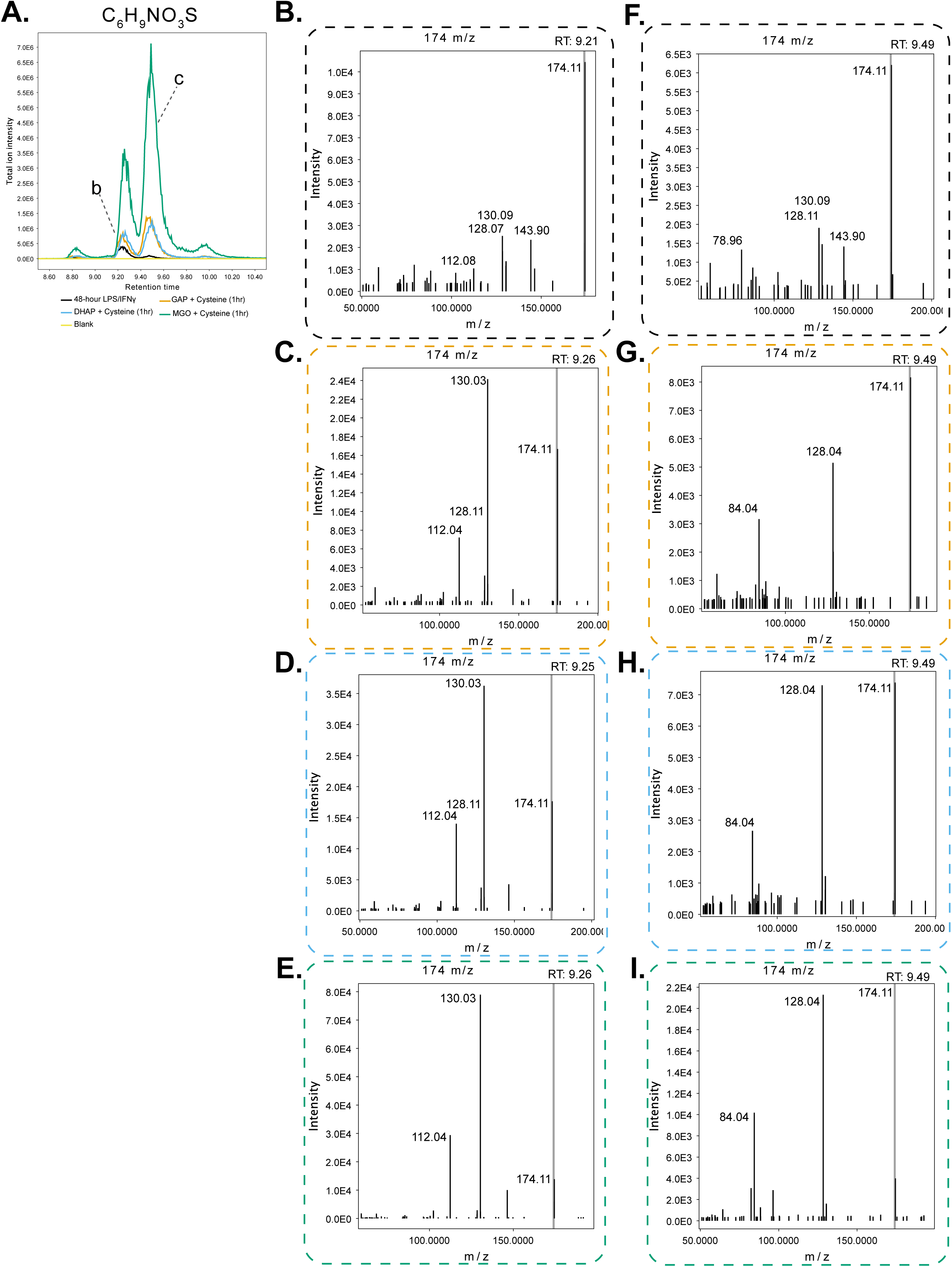
LC-MS/MS characterization of C_6_H_9_NO_3_S isomers. **A.** EIC at 174.023 m/z showing C_6_H_9_NO_3_S formation from DHAP (blue), GAP (yellow), or MGO (green) reacting with cysteine in vitro (1 hours), compared to LPS/IFNγ-stimulated BMDMs (black trace). **B-E.** MS/MS spectra of parent mass 174, at RT=9.2, in (B) cell extract of LPS/IFNγ-stimulated BMDMs, or the reaction mix of cysteine co-incubated with (C) GAP, (D) DHAP, or (E) MGO. This peak corresponds to the most abundant isomer of C_6_H_9_NO_3_S observed in cells (labeled as isomer b in Figure 1). (**F-I**) MS/MS spectra of parent mass 174, at RT=9.5, in (F) cell extract of LPS/IFNγ-stimulated BMDMs, or the reaction mix of cysteine co-incubated with (G) GAP, (H) DHAP, or (I) MGO. This peak corresponds to the second most abundant isomer of C_6_H_9_NO_3_S observed in cells (labeled as isomer c in Figure 1).

**Supplemental Figure 6.**
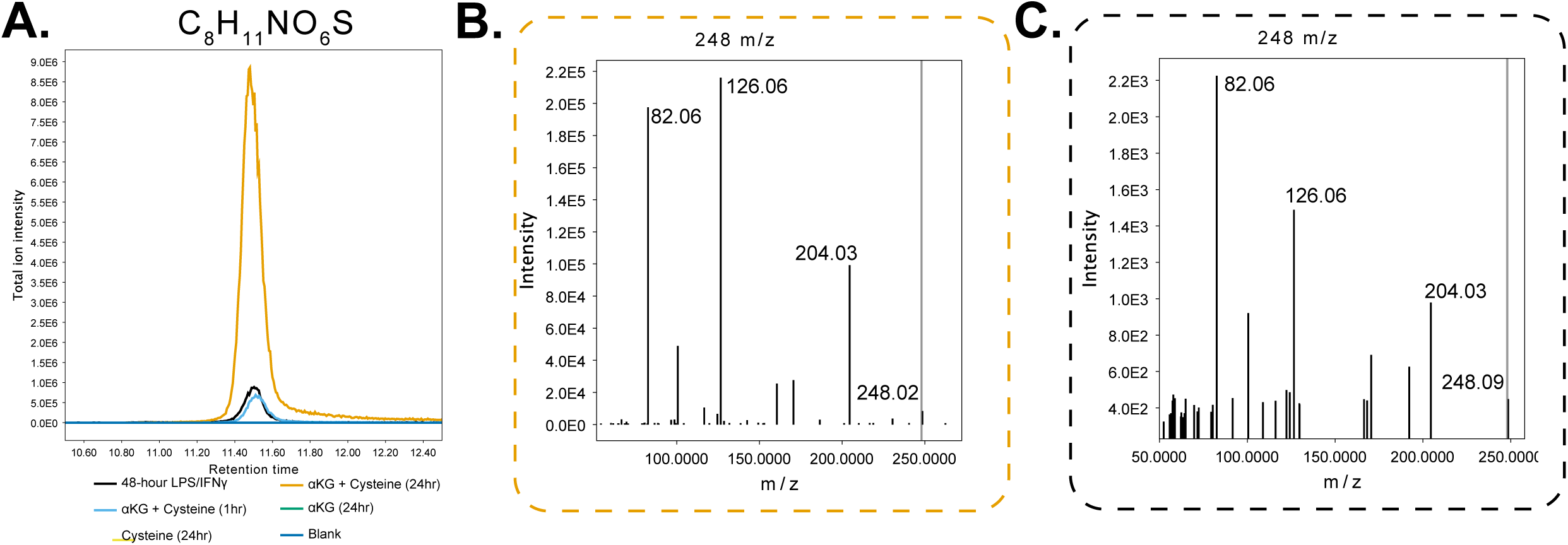
Chromatogram and MS/MS fragmentation patterns of αKG-Cys (C_8_H_11_NO_6_S) **A.** EIC at 248.023 m/z showing C_8_H_11_NO_6_S formation from αKG and cysteine in vitro co-incubation (1 and 24 hours) compared to LPS/IFNγ-stimulated BMDMs (black trace). **B-C.** MS/MS spectra correspond to the MS1 peak shown in A, in (**B**) reaction mix of αKG and cysteine in vitro co-incubation and (**C**) cell extract of LPS/IFNγ-stimulated BMDMs.

**Supplemental Figure 7.**
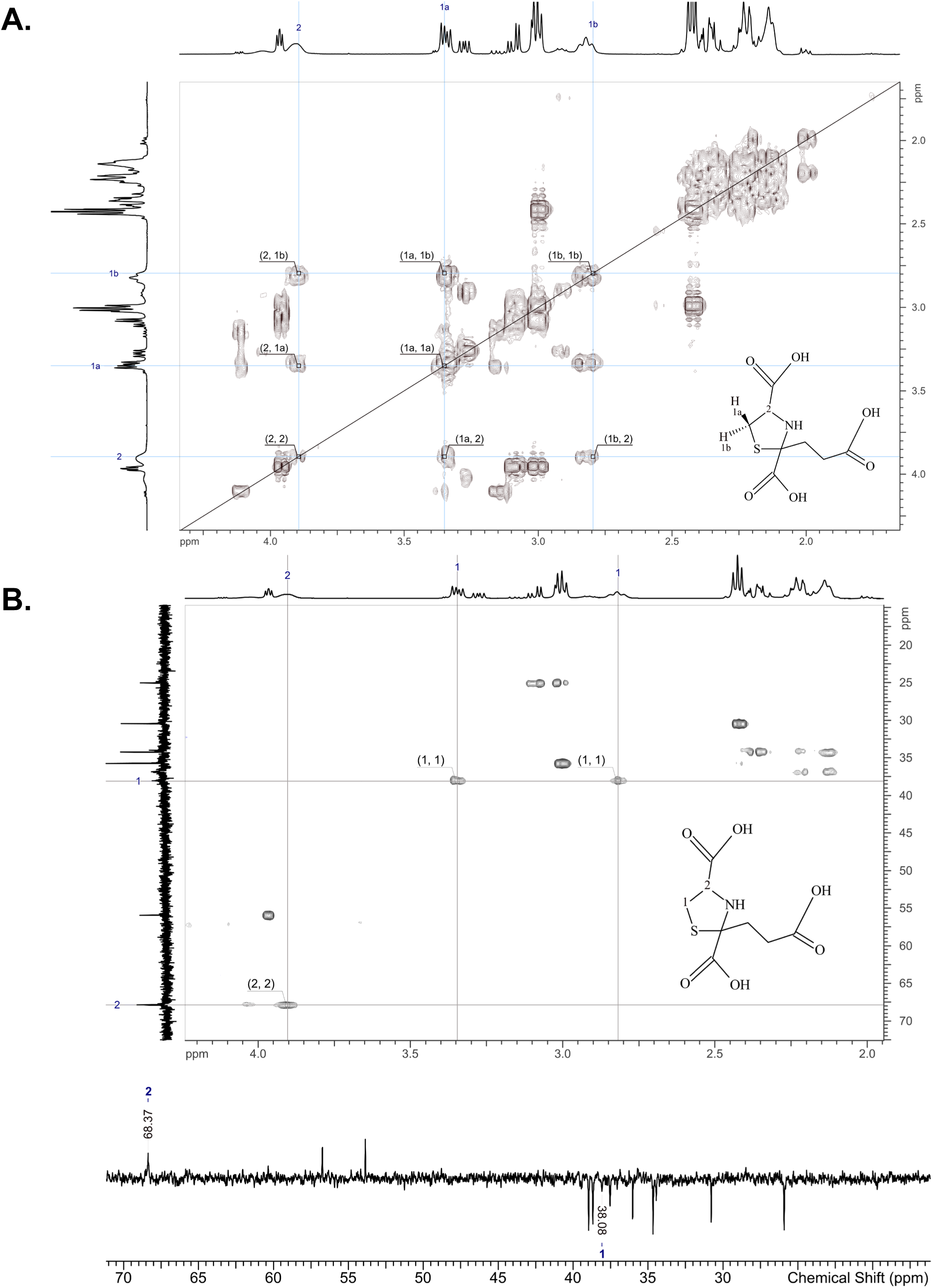
NMR structural analysis of αKG-Cys. **A.** ^1^H–^1^H COSY spectrum of the target compound showing the scalar-coupling network between protons 1a, 1b, and 2. The cross-peaks along the diagonal correspond to the individual proton resonances, while the off-diagonal correlations indicate through-bond coupling. Protons 1a and 1b exhibit a strong mutual cross-peak (1a↔1b), consistent with vicinal coupling within the same spin system. Both 1a and 1b also show clear correlations with proton 2 (1a↔2 and 1b↔2), confirming their connectivity. These COSY cross-peaks establish the proton–proton coupling, supporting the structural assignment indicated in the molecular diagram. No assignments were attempted in the 2.0–2.5 ppm region due to the high density of overlapping signals, which prevents meaningful or confident correlation analysis. **B.** 2D ^1^H–^13^C HSQC spectrum of the target compound, showing the direct proton–carbon correlations used to assign the spin system. The cross-peak associated with proton 1 aligns with the carbon resonance designated as carbon 1, establishing their direct connectivity. Likewise, the cross-peak associated with proton 2 correlates with carbon 2, confirming their assignments. The accompanying ^13^C DEPT-135 spectrum supports these assignments by identifying the carbon multiplicities and reinforcing the interpretation of the HSQC cross-peaks. Together, the HSQC and DEPT data provide consistent evidence for the structural assignments illustrated in the molecular diagram.

**Supplemental Figure 8.**
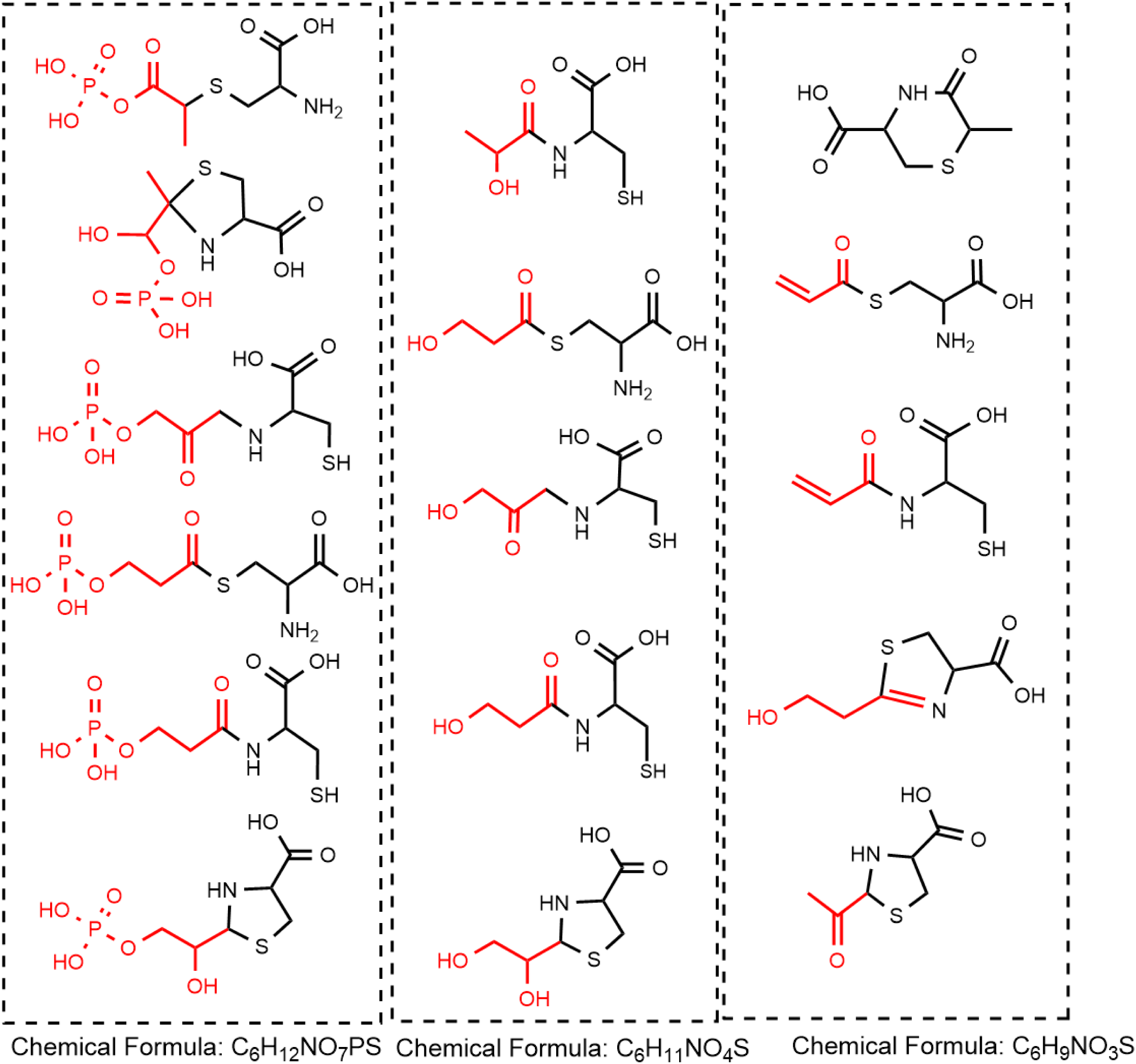
Schematic showing possible structures of cysteine adducts of formulas C_6_H_12_NO_7_PS, C_6_H_11_NO4S, and C_6_H_9_NO_3_S.

**Supplemental Figure 9.**
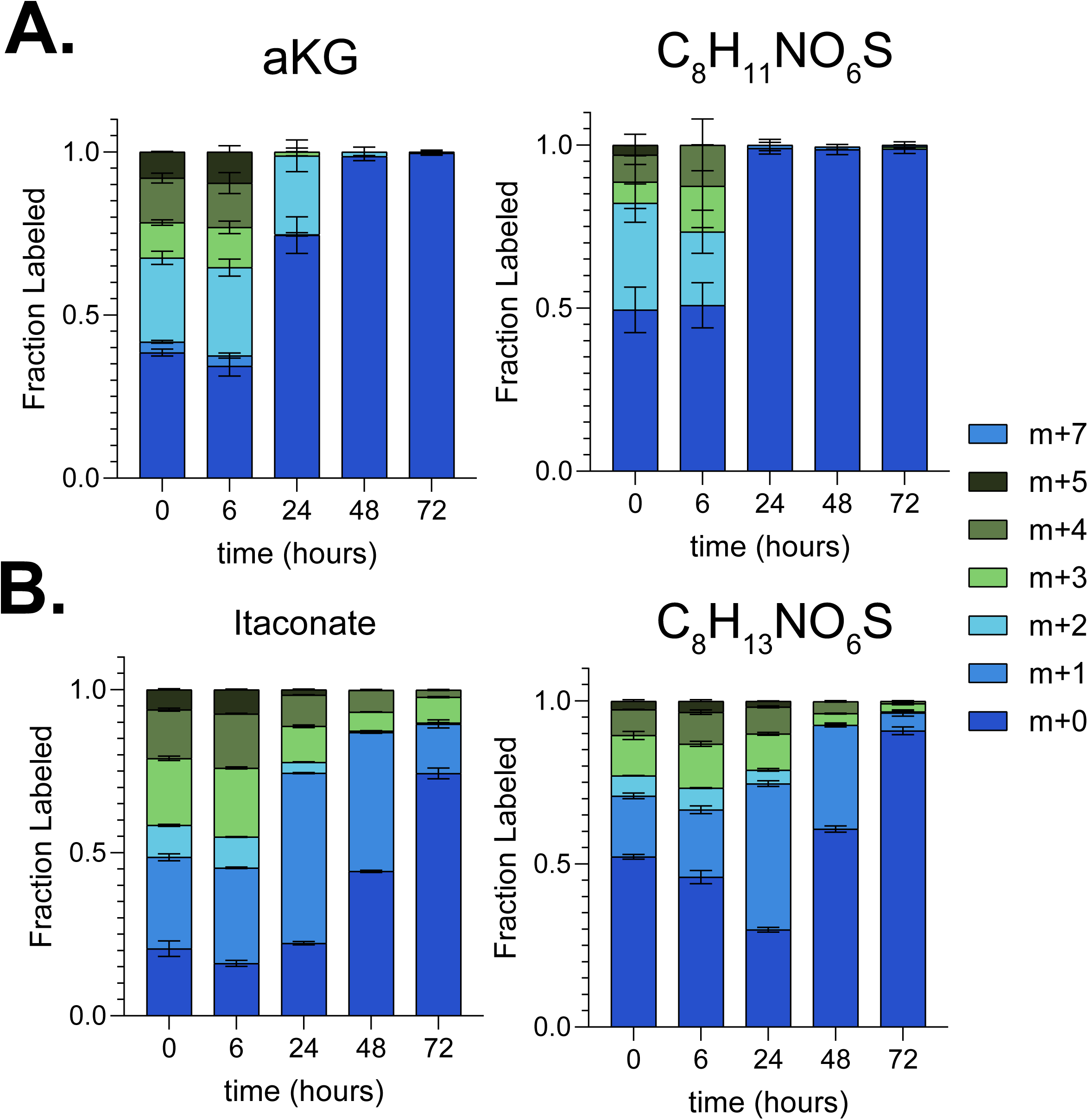
Isotope labeling pattern of TCA cycle-derived cysteine adducts over stimulation time course. **A.** Labeling patterns of αKG and αKG-Cys from ^13^C_6_-D-glucose in macrophages stimulated with LPS/IFNγ for indicated times. **B.** Labeling patterns of itaconate and Ita-Cys from ^13^C_6_-D-glucose in macrophages stimulated with LPS/IFNγ for indicated times. In all samples, cells were labeled in media containing ^13^C_6_-D-glucose tracer for 24h. Mean +/-standard deviation, n=3.

**Supplemental Figure 10.**
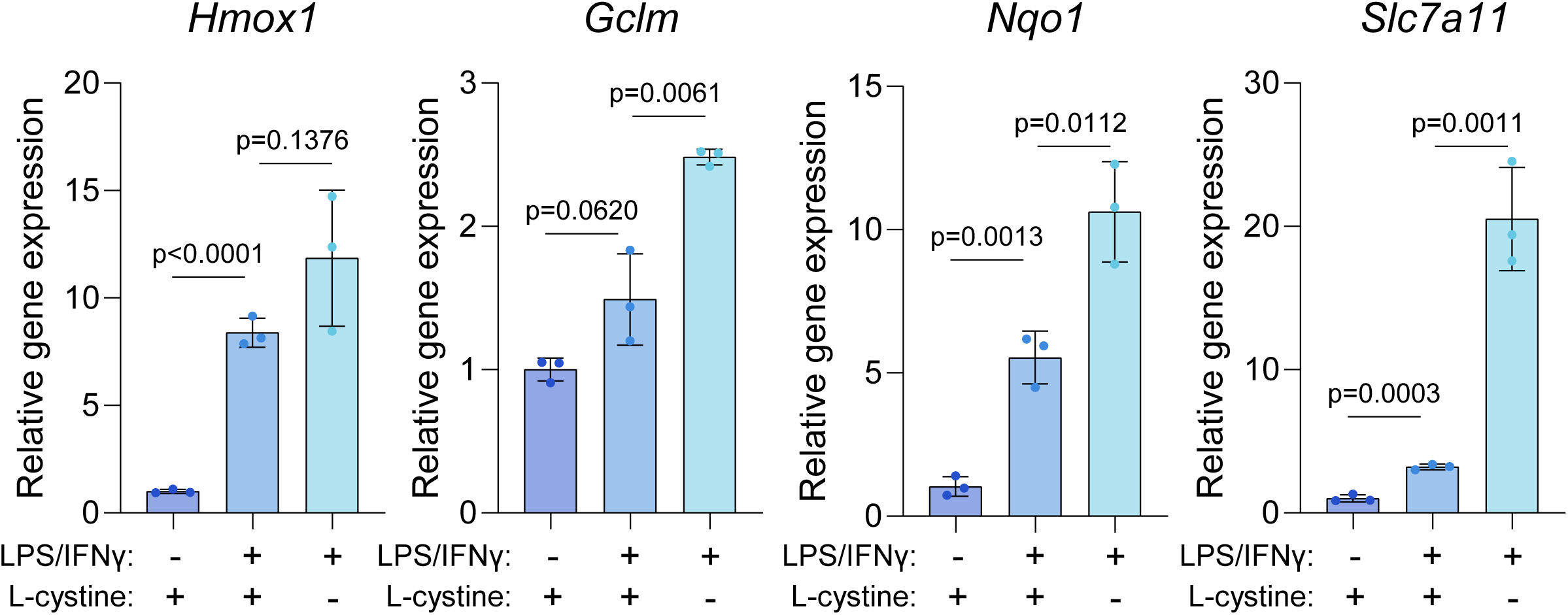
Classical activation and extracellular cystine availability modulate NRF2 targets. Relative mRNA expression of NRF2 target genes: *Hmox1*, *Gclm*, *Nqo1*, *Slc7a11* in BMDMs cultured in standard RPMI media (containing 200 μM cystine), with or without LPS/IFNγ-stimulation for 48 hours, or in LPS/IFNγ-stimulated BMDMs cultured in cystine deplete media. Mean +/- standard deviation, n=3. Statistical comparison by unpaired two-tailed t-tests with exact p-value reported.

